# A systematic analysis of diet-induced nephroprotection reveals overlapping and conserved changes in cysteine catabolism

**DOI:** 10.1101/2021.09.08.459468

**Authors:** Felix C. Koehler, Chun-Yu Fu, Martin R. Späth, K. Johanna R. Hoyer-Allo, Katrin Bohl, Heike Göbel, Jan-Wilm Lackmann, Franziska Grundmann, Thomas Osterholt, Claas Gloistein, Joachim D. Steiner, Adam Antebi, Thomas Benzing, Bernhard Schermer, Günter Schwarz, Volker Burst, Roman-Ulrich Müller

## Abstract

Caloric Restriction (CR) extends lifespan and augments cellular stress-resistance from yeast to primates, making CR an attractive strategy for organ protection in the clinic. Translation of CR to patients is complex, due to problems regarding adherence, feasibility and safety concerns in frail patients. Novel tailored dietary regimens, which modulate the dietary composition of macro- and micronutrients rather than reducing calorie intake promise similar protective effects and increased translatability. However, a direct head-to-head comparison to identify the most potent approach for organ protection as well as overlapping metabolic consequences has not been performed. We systematically analyzed six dietary preconditioning protocols - fasting mimicking diet (FMD), ketogenic diet (KD), dietary restriction of branched chained amino acids (BCAA), two dietary regimens restricting sulfur-containing amino acids (SR80/100) and CR - in a rodent model of renal ischemia-reperfusion injury (IRI) to quantify diet-induced resilience in kidneys. Of the administered diets, FMD, SR80/100 and CR efficiently protect from kidney damage after IRI. Interestingly, these approaches show overlapping changes in oxidative and hydrogen sulfide (H_2_S)-dependent cysteine catabolism as a potential common mechanism of organ protection. Importantly, these metabolic changes can be recapitulated in patients preconditioned by a diet limiting sulfur-containing amino acids indicating conserved diet-induced mechanisms of stress-resistance that may ultimately lead to clinical application.

## Introduction

Recent research has greatly improved our knowledge on longevity pathways and holds the promise to yield strategies that will allow for healthy aging in humans. Dietary protocols are among the most potent lifespan-extending interventions. Caloric Restriction (CR) - the simplest dietary longevity protocol - has been extensively studied and its effectiveness has been confirmed in a wide range of organisms ranging from budding yeast to rhesus monkeys (1, 2). This is a pivotal pre-requisite for potential translation to human medicine (3, 4). Mechanistically, the activation of longevity pathways leads to an increase in cellular and organismal stress-resistance (1, 5).

Regarding human disease, exploitation of this aspect is an important goal of translational research to protect organ function, when patients are exposed to damaging stimuli (e.g., surgery, ischemia, drug toxicity) in the clinical setting. In this context, the protection from acute kidney injury (AKI) is an important example. Being an aging-associated disease, the incidence of AKI has risen over the last decades (6). AKI is associated with increased mortality and morbidity and promotes progression to chronic kidney disease which, in turn, reduces both life expectancy and quality of life (7). Despite the considerable burden associated with AKI, both effective therapeutic and preventive measures are currently lacking. Interestingly, CR strongly protects from kidney damage in different rodent models of AKI (8–10). However, successful translation of dietary interventions into the clinical setting faces various obstacles partly due to problems regarding adherence, feasibility and safety concerns in frail patients. Consequently, developing tailored dietary interventions that are safe and feasible, but effective in high-risk patients is an important goal. Furthermore, an improved understanding of the molecular and metabolic mechanisms underlying the protective potential of different dietary regimens will be required to allow for the design of pharmacological strategies augmenting resilience.

Recent research has provided an insight into targeted dietary regimens, which - instead of reducing calorie intake - modulate the dietary composition of macro- and micronutrients (1, 11, 12). The fasting mimicking diet (FMD) mimics periodic fasting and enables the activation of cellular signal transduction similar to CR, such as a decrease in insulin, leptin and insulin growth factor 1 (IGF-1) (13–15). Furthermore, FMD reduces autoimmunity, inflammation and oxidative stress as well as it promotes autophagy resulting in cellular stress-resistance (11, 16). Ketogenic diets (KD) are high in fat and very low in carbohydrates and result in the synthesis of ketone bodies, such as β-hydroxybutyrate, exceeding β-oxidation of fatty acids as well as suppressing both inflammation and oxidative stress (17, 18). Considering that CR in rodents actually induces ketogenesis due to the resulting feeding cycles and fasting itself, the examination of the protective properties of ketogenic diet are warranted (19). Dietary inhibition of the mechanistic Target of Rapamycin (mTOR) pathway is achieved when the branched chain amino acids (BCAA) valine, leucine and isoleucine are limited (12, 20). mTOR inhibition is known to result in augmented cellular stress-resistance by reducing protein translation and decreasing proteotoxic and oxidative stress (21, 22). In addition, restriction of the sulfur-containing amino acids (SAA) methionine and cysteine is an attractive target to mimic the nephroprotective effect of CR (23). Depletion of methionine and/or cysteine has been widely studied in various longevity model organisms and can be achieved by consuming SAA free or low-SAA foods such as fruit, refined grains and vegetables in humans (12, 24). Reduction of SAA leads to an increase in lifespan and improves metabolic health observed across taxa (1). The balance between methionine and cysteine is regulated by the trans-sulfuration pathway (TSP) and SAA restriction is associated with anti-inflammation, increased stress-resistance, reduction of reactive oxygen species, decline in insulin signaling and alteration in lipid-metabolism (12, 25, 26). All the described dietary approaches have been studied with respect to feasibility and safety in clinical trials or are already in clinical use (14, 27–31). Thus, their administration as a preconditioning strategy in the clinical setting could add substantially to our therapeutic armamentarium. However, systematic analyses on the power and efficacy of different dietary approaches are limited and a direct comparison of these protocols is lacking. While CR has been studied extensively in rodent models of kidney injury, FMD, KD and BCAA restriction have not been examined in this regard (8–10). Addition of SAA to protein-restricted dietary regimens has been shown to abrogate the protective potential, however, SAA restriction itself has not been tested to date (23). Besides, underlying metabolic patterns of protective strategies have not been compared and knowledge on their conservation in humans is limited.

Consequently, we set out to perform a simultaneous comparison of six dietary interventions (FMD, KD, BCAA, SR80/100 - two dietary regimens either reducing sulfur-containing amino acids (SAA) by 80 percent (SR80) or fully depleting them (SR100) and CR) in a murine model of AKI. Besides, profiling of metabolites and proteomics were employed to provide an insight into the molecular mechanisms underlying dietary organ protection. Analyses in a human cohort were used to confirm evolutionary conservation of the metabolic response.

## Results

### Weight development and baseline kidney function in dietary preconditioned and non-preconditioned mice

To examine the diet-induced changes in phenotype and functional kidney parameters, mice were treated with the different dietary regimens (CR, FMD, the low-SAA diets SR80/100, KD and BCAA) for 14 days, sacrificed and compared to *ad libitum* fed, non-preconditioned (non-PC) animals. CR and SR100 mice lost weight significantly, whereas KD mice gained weight compared to non-PC mice. Treatment with FMD, SR80 or BCAA did not result in any weight differences (Fig. S1A). Renal function was assessed by measuring serum creatinine and serum urea and there were no differences in creatinine across all dietary regimens. CR - in line with previously published data - led to a significant increase and KD to a significant decrease in serum urea compared to the non-PC group after 14 days of exposure to the dietary regimens (9). FMD, SR80/100 and BCAA did not reveal significant changes in serum urea (Fig. S1B, C). Additionally, none of the diets had an impact on kidney tissue morphology as determined through a blinded analysis of periodic acid–Schiff (PAS)-stained paraffin-embedded tissue (Fig. S1D, E) (32, 33). Furthermore, we did not observe any diet-related changes of general health, behavior or activity of the mice.

### Tailored dietary preconditioning improves survival and general health status in mice after renal IRI

To induce AKI, we performed right nephrectomy and exposed the left kidney to IRI in male C57BL/6N mice after dietary preconditioning and respective controls. CR and the non-PC group served as positive and negative controls, respectively. Sham animals, which underwent right nephrectomy without clamping of the vascular pedicle of the contralateral kidney, did not develop AKI and served as an additional control for the surgical setting (Fig. 1A). Preconditioning with FMD, SR80/100 and CR led to significantly improved survival following AKI (*P < 0.0001*). None of these animals died or had to be euthanized according to previously defined criteria within 72 hours after renal IRI, whilst this was the case for 50% of the non-PC animals. Two out of 13 (15.4%) and nine out of 13 (69.2%) animals had to be sacrificed or died within 72 hours after renal IRI in the KD and the BCAA group, respectively (Fig. 1B). Moreover, animals in the SR100 and the CR group gained weight within 72h, whereas FMD, SR80, KD, BCAA, non-PC and even sham animals lost weight after surgery (Fig. 1C). In line with the survival benefit, mice preconditioned with FMD, SR80/100, KD or CR showed an improved overall health status both 24(Table S1, Fig. S2).

**Fig. 1.**
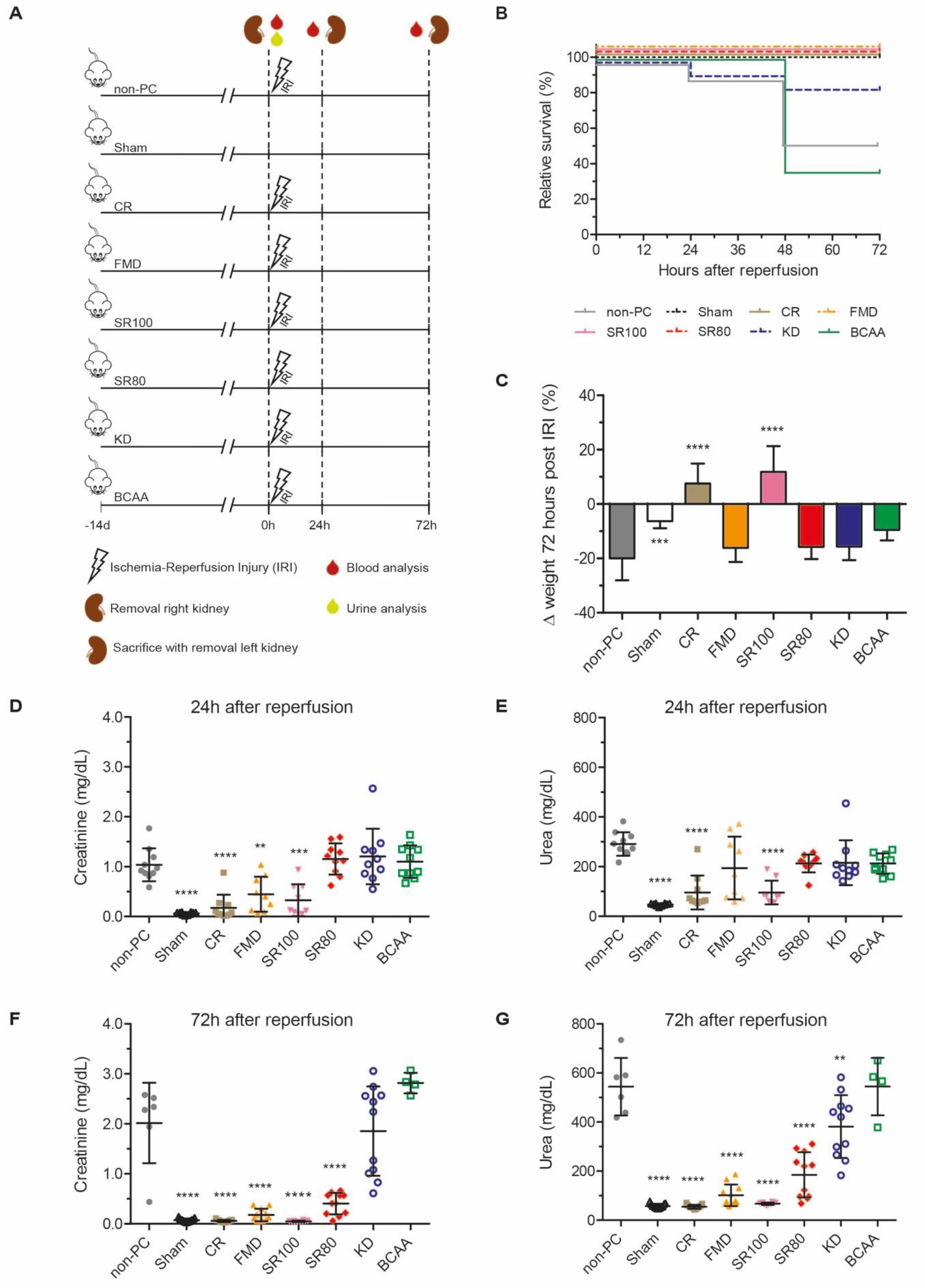
Direct comparison of different tailored dietary interventions identifies FMD and SR80/100 as potent approaches protecting from renal IRI beyond CR. (A) Schematic of the experimental design depicting dietary preconditioning modes. CR served as positive control, non-PC as negative. Sham animals underwent right nephrectomy without clamping of the left renal pedicle and do not develop acute kidney injury. (B) Survival curves of dietary preconditioned, non-preconditioned and sham animals after renal IRI. FMD, KD and both low-SAA diets SR80/100 led to a significant survival advantage. (C) Delta body weight of dietary preconditioned, non-preconditioned and sham animals 72 hours after reperfusion or surgery, respectively, compared to the respective animal before IRI. Data are represented as mean ± standard deviation. (D, E) Creatinine and urea concentrations 24 hours after reperfusion or surgery, respectively. (F, G) Creatinine and urea concentrations 72 hours after reperfusion or surgery, respectively. Kaplan Meier Analysis (log rank test *\*\*\*\*P < 0.0001*, n = 13 per group) was used in (A), One-way ANOVA and Bonferroni multiple comparison test across all dietary preconditioning groups indicating differences to non-PC mice (*\*\*P < 0.01*, *\*\*\*P < 0.001* and *\*\*\*\*P < 0.0001*) was used in (C)-(G) Data are represented as mean ± standard deviation in (C)-(G) and each mice is depicted by a single dot. Abbreviations: BCAA: Dietary Restriction of Branched Chain Amino Acids; CR: Caloric Restriction; d: day; FMD: Fasting Mimicking Diet; h: hour; IRI: Ischemia-Reperfusion Injury; KD: Ketogenic Diet; non-PC: non-preconditioned animals; SR100: 100% Restriction of Sulfur-containing Amino Acids; SR80: 80% Reduction of Sulfur-containing Amino Acids.

### FMD and SR80/100 recapitulate CR benefits in maintaining kidney function following IRI

Kidney damage of the different groups was quantified by measuring serum creatinine and serum urea at 24 hours and 72 hours after renal IRI. At 24 hours, FMD, SR100 and CR showed significantly lower creatinine values compared to the non-PC group, whilst creatinine did not differ between non-PC, SR80, KD and BCAA (Fig. 1D). Regarding serum urea, preconditioning with SR100 and CR significantly attenuated the increase observed at 24 hours after IRI, whereas FMD, SR80, KD and BCAA did not lead to a significant difference in serum urea compared to the non-PC group (Fig. 1E). With regard to the later time-point 72h after IRI, in contrast to non-PC animals, kidney function returned to near normal in FMD, SR80/100 and CR mice. Regarding mice on KD, only serum urea levels were significantly lower than in non-PC animals. With respect to serum creatinine levels, KD preconditioned mice apparently split into two groups. BCAA preconditioning did not result in any signs of renal protection (Fig. 1F, 1G). The sham intervention did not have any impact on kidney function (Fig. 1 D-G).

When compared to baseline values obtained from mice that were sacrificed after the preconditioning phase (i.e., before surgery), creatinine concentrations were not significantly elevated at 24 hours and 72 hours after IRI in CR or SR100 preconditioned mice (Fig. S3A). SR100 preconditioned mice showed slightly elevated serum urea values at 24 hours, whilst there was no significant rise at 72 hours (Fig. S3B). CR mice did not differ significantly in serum urea at 24 hours, however, CR resulted in elevated values already at baseline (Fig. S1C). FMD and SR80 preconditioned mice revealed mildly impaired kidney function at 24 hours after renal IRI as indicated by an increase in both serum creatinine and urea at this time-point compared to respective baseline animals. However, at 72 hours after IRI, kidney function was almost completely restored in FMD and SR80 animals (in SR80 serum urea remained slightly elevated). In contrast, KD- and BCAA-preconditioned, as well as non-PC mice revealed a severely impaired kidney function at all examined points in time compared to respective baselines animals (Fig. S3).

### FMD, SR80/100 and CR strongly attenuate cellular damage after renal IRI

To assess damage in epithelial cells of the renal tubular system after IRI, we used an established semi-quantitative histologic damage score integrating epithelial flattening, loss of brush borders, loss of nuclei, necrosis and vacuolization determined in a blinded manner (32, 33). Direct comparison of the overall damage score identified significantly less acute tubular damage in FMD, SR100 and CR preconditioned mice compared to non-PC animals 24 hours after IRI (Fig. S4A). This protection from kidney damage was mainly driven by the extent of tubular necrosis. Differences in other common signs of tubular damage did not reach statistical significance at this early time-point (Fig. S4B-G). Neither the overall score nor necrosis quantified in KD and BCAA animals differed from non-PC mice 24S4A, S4F, S4GFMD, the low-SAA diets SR80/100 and CR mice sacrificed at 72 hours after IRI were significantly protected from renal damage quantified by histology (Fig. 2A). Again, this result was primarily caused by differences in the extent of tubular necrosis (Fig. 2B, S5A-S5D). However, SR100 animals additionally revealed significantly less loss of brush borders compared to non-PC mice at this time-point (Fig. S5C). To perform a longitudinal analysis after renal IRI within each group, damaged kidneys obtained at 24 hours and 72 hours after IRI were also compared to baseline animals of the respective dietary group confirming the protective effects. In contrast, in KD, BCAA and non-PC mice acute tubular damage significantly worsened over time after renal IRI, both when compared to non-PC kidneys or the respective baseline tissue (Fig. 2, S4, S5). To further analyze the degree of cell death, we performed immunostaining of cleaved-caspase-3 and deoxynucleotidyl transferase–mediated dUTP nick end-labeling assay (TUNEL) (34, 35). Immunostaining of cleaved-caspase-3 suggested severe cellular damage in non-PC and BCAA mice, whereas KD showed a less profound staining. FMD, the low-SAA diets SR80/100 and CR animals showed (almost) no anti-cleaved-caspase-3 positive staining pattern (Fig. S6). TUNEL staining demonstrated high numbers of positive cells in BCAA and non-PC animals, significantly less in KD and SR80 animals and hardly any signal in FMD, SR100 and CR animals (Fig. 2D).

**Fig. 2.**
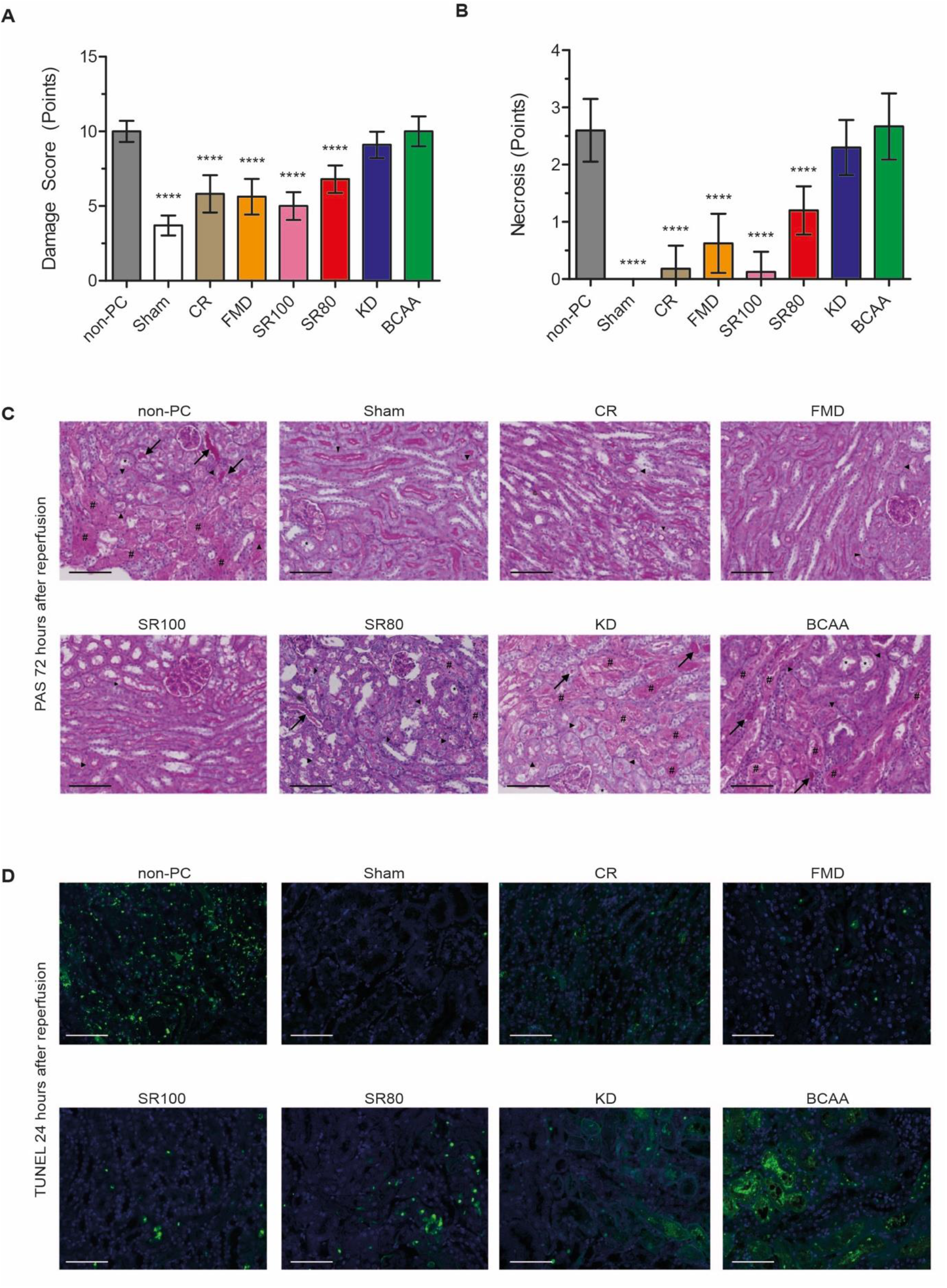
Dietary preconditioning by FMD, SR80/100 and CR efficiently prevent tissue damage and cell death after renal IRI. **(A, B)** Histologic damage and necrosis score at 72 hours after reperfusion examined in dietary preconditioned, non-preconditioned and sham animals. One-way ANOVA and Bonferroni multiple comparison test across all dietary preconditioning groups indicating differences to non-PC mice (*\*\*\*\*P < 0.0001*). Data are represented as mean ± standard deviation. 24h after IRI: n = 10 per group; 72h after IRI n = 13 in the FMD, SR80/100, CR and sham group, n = 9 in the KD, n = 4 in the BCAA and n = 6 in the non-PC group, respectively. hours after reperfusion Arrowheads indicate nuclei loss, arrows tubular casts, * epithelial flattening and # denuded tubuli with luminar debris, respectively.Terminal deoxynucleotidyl transferase– mediated digoxigenin-deoxyuridine nick-end labeling (TUNEL) assay at 24 hours after reperfusion from dietary preconditioned, non-preconditioned and sham animals. Cell death in green. Abbreviations: BCAA: Dietary Restriction of Branched Chain Amino Acids; CR: Caloric Restriction; FMD: Fasting Mimicking Diet; KD: Ketogenic Diet; non-PC: non-preconditioned animals; PAS: SR100: 100% Restriction of Sulfur-containing Amino Acids; SR80: 80% Reduction of Sulfur-containing Amino Acids; TUNEL: Terminal deoxynucleotidyl transferase–mediated digoxigenin-deoxyuridine nick-end labeling.

### FMD, SR80/100 and CR modulate oxidative cysteine and hydrogen sulfide metabolism with a central role for sulfite

CR promotes the H_2_S-generating branch of cysteine metabolism (TSP) resulting in the generation of H_2_S, which has been shown to be a mediator of CR-induced stress resistance (23). Supplementation of sulfur-containing amino acids counteracts the CR-orchestrated benefits in rodents (23). Based on this knowledge, we inspected the four most potent dietary regimens (FMD, SR80/100 and CR) regarding their impact on cysteine metabolism in post-diet pre-IRI mice. Targeted quantification by high performance liquid chromatography (HPLC) revealed a significant elevation of serum cysteine after FMD and, surprisingly, also in the cysteine-free diet SR100. The increase observed in the SR80 group did not reach statistical significance and CR did not change serum cysteine compared to the *ad libitum* fed non-PC group (Fig. 3A). In contrast, serum homocysteine was commonly reduced at baseline after FMD, the low-SAA diets and CR compared to non-PC animals indicating cysteine turnover (Fig. 3B).

**Fig. 3.**
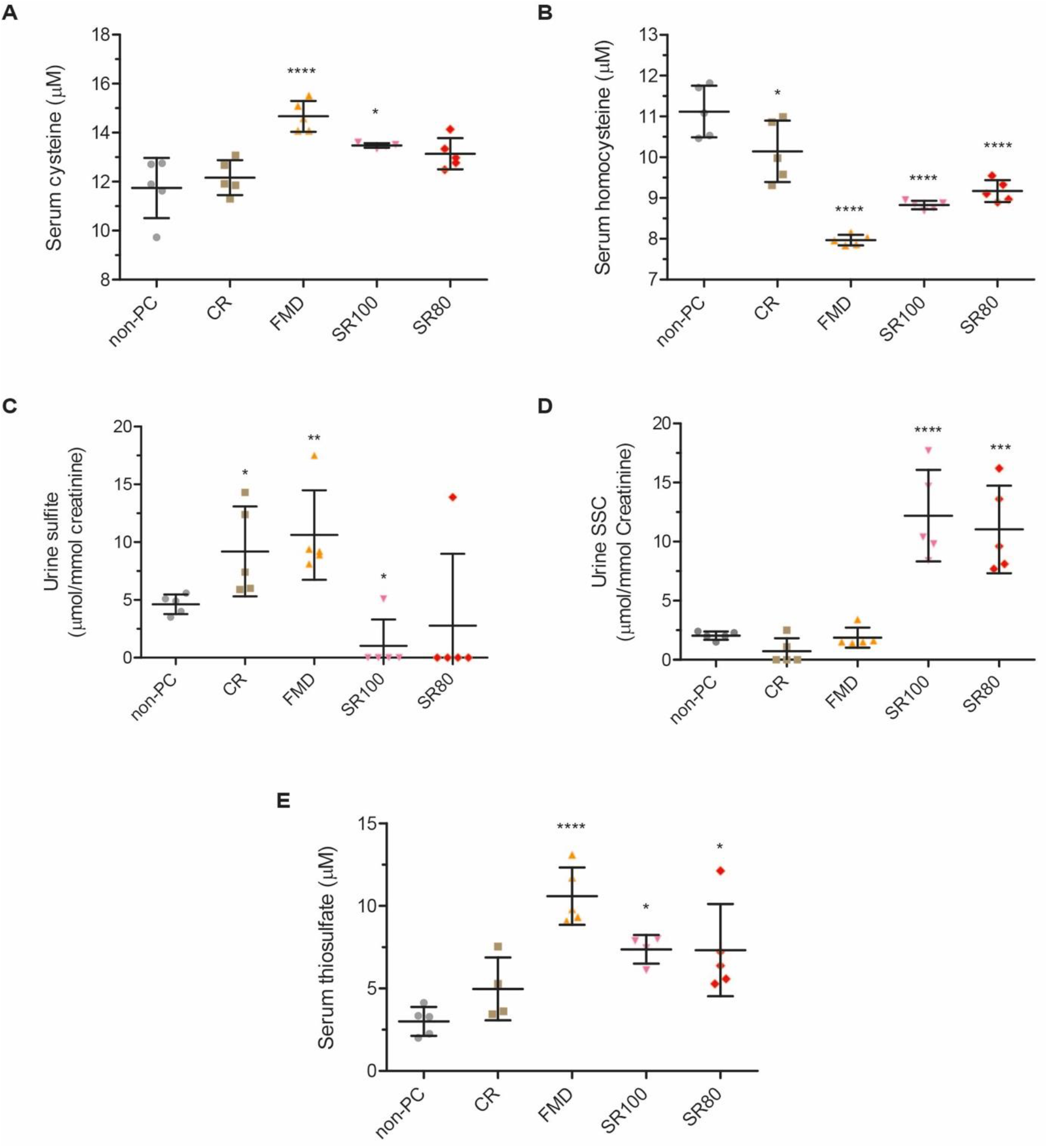
FMD, SR80/100 and CR modulate the oxidative and hydrogen sulfide dependent cysteine catabolism with sulfite as key intermediate. **(A, B)** Serum cysteine and homocysteine values measured in dietary preconditioned and non-preconditioned mice at baseline. **(C)** Urine sulfite values measured in dietary preconditioned and non-preconditioned mice at baseline. **(D)** Urine s-sulfocysteine (SSC) values measured in dietary preconditioned and non-preconditioned mice at baseline. **(E)** Serum thiosulfate values measured in dietary preconditioned and non-preconditioned mice at baseline. One-way ANOVA and Bonferroni multiple comparison test across all dietary preconditioning groups indicating differences to non-PC mice (\**P <0.05*, *\*\*P < 0.01*, *\*\*\*P < 0.001* and *\*\*\*\*P < 0.0001*) was used in (A), (B), (D) and (E). Unpaired T-test between dietary preconditioned and non-preconditioned mice (*\*P < 0.05* and *\*\*P < 0.01*) was used in (C). All data are represented as mean ± standard deviation and each mice is depicted by a single dot. Abbreviations: CR: Caloric Restriction, FMD: Fasting Mimicking Diet; non-PC: non-preconditioned mice; SR100: 100% Restriction of Sulfur-containing Amino Acids; SR80: 80% Reduction of Sulfur-containing Amino Acids, SSC: S-sulfocysteine.

Next, we characterized metabolites of cysteine catabolism in more detail by quantifying s-sulfocysteine (SSC), thiosulfate and sulfite levels. SSC and thiosulfate are two major end-products of the cysteine catabolism, whilst sulfite serves as a common intermediate of both catabolic pathways and precursor of SSC (Fig. 4). Urine sulfite measurements revealed significantly elevated levels in both FMD and CR as compared to non-PC animals. In contrast, we found a decrease in urine sulfite in the low-SAA diets (Fig. 3C). Of note, SR80/100 resulted in a profound rise in urine SSC as compared to non-PC animals with no significant difference between the two SAA restricted regimens, an effect that was neither induced by FMD nor CR (Fig. 3d). SSC is generated by the non-catalyzed reaction of cystine and sulfite and is widely used as a biomarker for sulfite levels indicating elevated sulfite signaling in low-SAA diets. Increased turnover of sulfite towards cysteine and SSC may explain both the reduced urine sulfite levels and the lack of a reduction (SR80) or even rise (SR100) in serum cysteine observed upon dietary SAA restriction (Fig. S7).

**Fig. 4.**
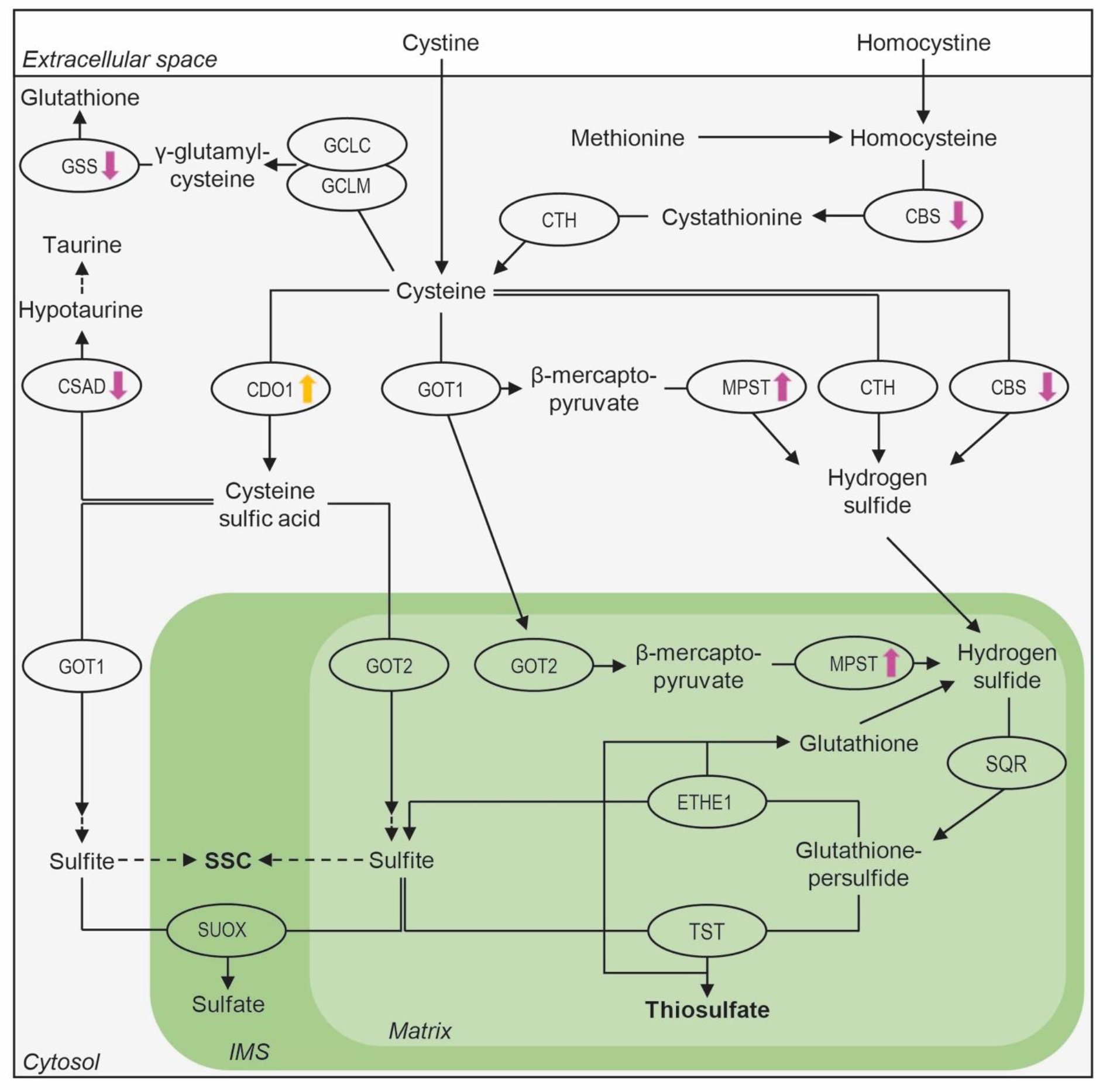
Key enzymes in cysteine catabolism are differentially regulated in response to FMD and SR100. Cysteine is either imported or generated from methionine and cysteine is either degraded to glutathione or taurine, or it is metabolized via the oxidative or the hydrogen sulfide (H_2_S)-dependent branches to sulfate, thiosulfate and s-sulfocysteine (SSC), respectively. Sulfite is the major intermediate in both oxidative and H_2_S-dependent cysteine catabolism. Enzymes significantly regulated *(p<0.05, log2FC > 0.6 or < −0.6*) after FMD or SR100 are assigned by colored arrows (Fasting Mimicking Diet (FMD): orange; 100% Restriction of Sulfur-containing Amino Acids (SR100): pink). Enzymes are depicted as ellipsis. Arrows with solid line indicate enzymatic, arrows with dashed lines non-enzymatic reactions, respectively. The investigated major end-products of the H_2_S-generating branch SSC and thiosulfate are written in bold. Abbreviation: IMS: mitochondrial intermembrane space; SSC: S-sulfocysteine.

Thiosulfate - another end-product in H_2_S-dependent cysteine metabolism - is generated by H_2_S catabolism in a sulfite-dependent manner. We found significantly increased serum thiosulfate levels after FMD and SR80/100 as well as a strong trend towards elevated serum thiosulfate levels in CR animals (*P = 0.08*) compared to non-PC animals (Fig. 3e). Conclusively, FMD, low-SAA diets and CR result in a commonly shared activation of both oxidative and H_2_S-dependent cysteine catabolism as demonstrated by elevated thiosulfate, sulfite and SSC (Fig. S7).

### SR100 results in a reciprocal regulation of H_2_S-synthesizing enzymes MPST and CBS

We used quantitative proteome profiling to decipher changes of key enzymes in the TSP in kidney in response to SAA depletion and FMD (Fig. 4, S7, S8, Table S2). Interestingly, SR100 - but none of the other regimens - lead to upregulation of MPST that catalyzes the desulfuration of 3-mercaptopyruvate (3-MP) and transfers sulfane sulfur to its thiophilic acceptors resulting in release of H_2_S. This increase in MPST, was accompanied by a significant decrease of CBS levels in response to low SAA levels, whereas there was no significant change in CTH, the known TSP enzymes involved in H_2_S biosynthesis.

Independent from the H_2_S-generating branch, cysteine is metabolized via two other distinct pathways, the γ-glutamylcysteinylglycine dependent synthesis of glutathione and the branch leading to the formation of either taurine or sulfate. Of interest, the key enzymes in both glutathione, as well as taurine synthesis, GSS and CSAD, respectively, were significantly downregulated in our animals in response to SR100 (Fig. 4, Table S2). Taken together, dietary SAA depletion results in a profound modulation of key enzymes within cysteine metabolism in favor of MPST-dependent activation of the H_2_S-generating branch in murine kidneys.

### FMD leads to an upregulation of CDO1 and higher expression levels of GOT1/2 in oxidative and H_2_S-dependent cysteine catabolism

CDO1 is the first and rate-limiting enzyme in oxidative cysteine catabolism and converts cysteine to cysteine sulfic acid (CSA). FMD resulted in a strong upregulation of CDO1 (Table S2). CDO1 is tightly regulated in cysteine homeostasis and elevated levels of CDO1 correlate with increased cysteine abundance accompanied by an activation of oxidative cysteine catabolism. In line with the marked increase in CDO1, the expression levels of the downstream targets GOT1/2 were also elevated in response to FMD; however, this upregulation was less strong and less significant (Fig. S8, Table S2). GOT1 has been shown recently to function in CSA deamination leading to sulfite, while GOT2 is part of an MPST-dependent pathway resulting in the generation of H_2_S (Fig. 4) (36).

### Proteome profiling identifies overlapping regulation of autophagy associated proteins in beneficial diets

To decipher overlapping diet-mediated molecular changes beyond cysteine and H_2_S metabolism, we performed a global quantitative analysis of the proteome data. Principal component analysis revealed the most profound separation for FMD, SR100 and CR animals from non-PC mice, whereas kidneys after preconditioning with SR80, KD and BCAA tended to be more similar (Fig. S9A). To identify common patterns in dietary regimens that were clearly associated with renal organ protection we compared FMD, SR80/100 and CR in more detail. These protocols induced differential regulation of 176, 87, 377 and 132 proteins compared to non-PC animals (Fig. S9B). A significant overlap was observed in 48 proteins that were differentially expressed in kidneys in response to at least three of these dietary strategies. Seven proteins (ATG7, BCR, CTPS2, KCNJ16, PRKA, SPATA5, TMEM256) revealed a shared mode of regulation after dietary preconditioning with FMD, SR80/100 and CR compared to non-PC animals (Fig. S9C). Of note, three commonly regulated proteins (ATG7, CTPS2, PRKA) are associated with cellular processes in autophagy.

### Generation of SSC and elevated sulfite metabolism are induced by a low-SAA diet in humans

To confirm whether molecular mechanisms obtained by low-SAA diets are conserved in humans, changes in cysteine catabolism were analyzed in a pilot-cohort of ten participants that underwent dietary SAA restriction using a formula diet for seven days compared to ten age- and sex-matched individuals on a formula diet containing standard levels of SAA (Table S3).

To decipher changes in cysteine catabolism upon dietary SAA restriction in humans we quantified serum cysteine, serum cystine und urinary SSC levels in the low-SAA and the control group. In line with our findings from the rodent studies, pre- and post-diet cysteine levels, as well as the individual turnover in cysteine did not differ between the two groups even though SAA were heavily restricted (Fig. 5A-C). As cystine reacts with sulfite to generate SSC and cysteine, we detected significant lower levels of serum cystine in response to the low-SAA group, however, the individual fold change did not reach statistical significance (Fig. 5D-F) (37). Urinary SSC levels, a major side-product of sulfite elevation, did not differ between the low-SAA and the control group at baseline prior to dietary intervention (Fig. 5G, 5J). Of interest and consistent with our rodent data, participants in our pilot-cohort receiving the SAA-restricted diet had significantly higher urinary SSC levels in both spontaneous samples and 24h urine collections after short term dietary intervention (Fig. 5H, 5K). Besides, the individual increase of urinary SSC was significantly higher in the low-SAA group (Fig. 5I, 5L). Considering the significant findings regarding SSC and its central role in the H_2_S-producing branch of cysteine catabolism, we went on to measure SSC in a larger group of patients (n=49) after SAA restriction. Again, pre-dietary levels of urinary SSC did not differ (Fig. 6A) and we observed a significant rise in urinary SSC in response to the low-SAA diet indicating active cysteine catabolism (Fig. 6B, 6C).

**Fig. 5.**
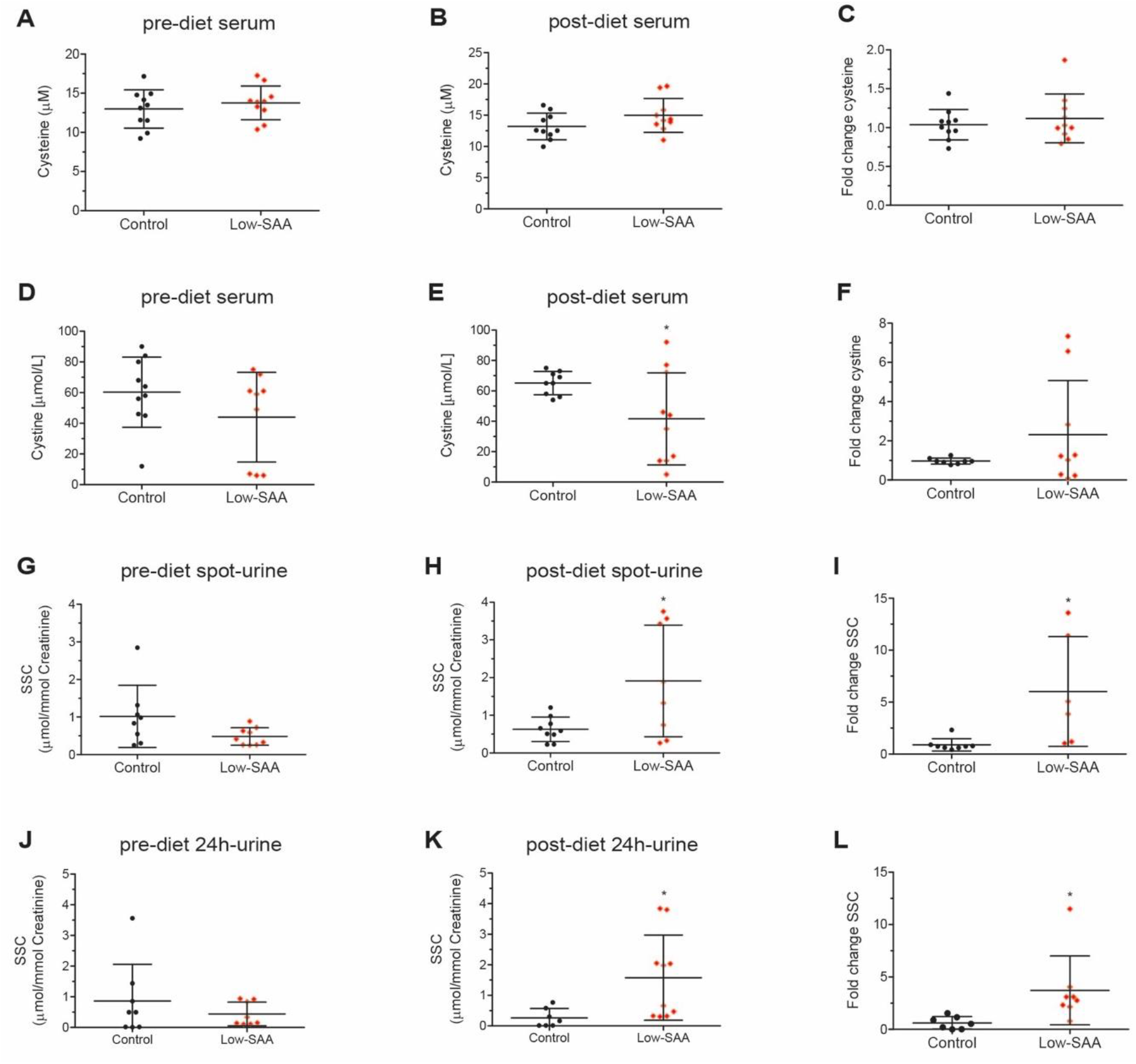
Metabolic changes in oxidative and hydrogen sulfide (H_2_S)-dependent cysteine catabolism are induced by short-term dietary preconditioning with low-SAA in humans. **(A, B)** Cysteine concentrations measured in human serum samples after 7 days of control or low-SAA diet, before and after dietary intervention. **(C)** Individual fold change of cysteine levels in serum of the control and low-SAA groups. **(D, E)** Cystine concentrations measured in human serum samples after 7 days of control or low-SAA diet, before and after dietary intervention. **(F)** Individual fold change of serum cystine levels in the control and low-SAA groups. **(G, H)** S-sulfocysteine (SSC) concentrations measured in human spontaneous urine (spot-urine) samples after 7 days of control or low-SAA diet, before and after dietary intervention. **(I)** Individual fold change of SSC levels in spot-urine in the control and low-SAA groups. **(J, K)** SSC concentrations measured in 24h collected human urine samples after 7 days of control or low-SAA diet, before and after dietary intervention. **(L)** Individual fold change of SSC levels in 24h collected urine samples in the control and low-SAA groups. Unpaired T-test of low-SAA compared to control diet (*\*P < 0.05*) was used in (A)-(L). All data are represented as mean ± standard deviation and each participant is depicted by a single dot. Abbreviations: h: hour; SAA: Sulfur-containing Amino Acids; spot-urine: spontaneous urine; SSC: S-sulfocysteine.

**Fig. 6.**
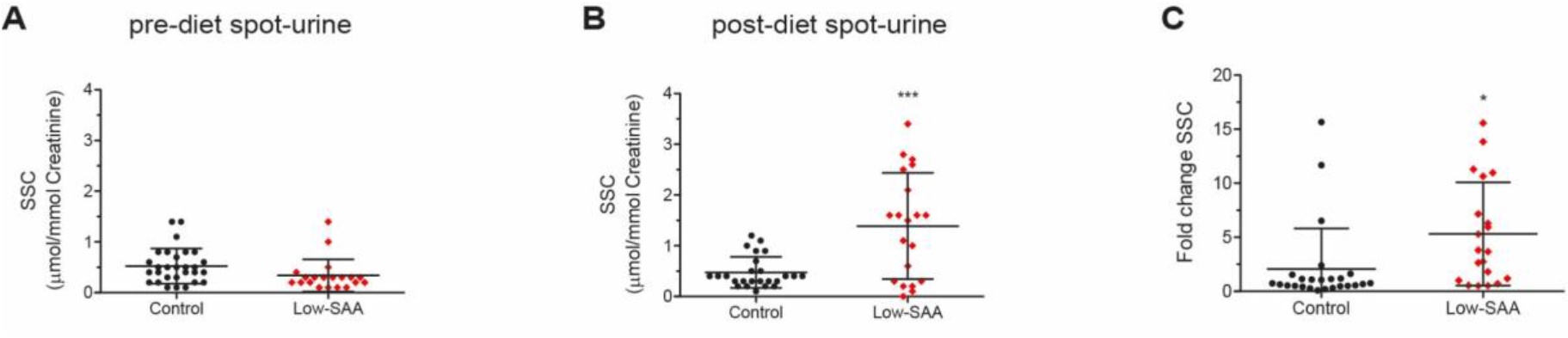
S-sulfocysteine – a key end-product of the H_2_S generating branch of cysteine catabolism – is induced upon a low-SAA diet in humans. **(A, B)** S-sulfocysteine (SSC) concentrations measured in spontaneous urine (spot-urine) samples of patients before and after the dietary intervention (control (N=29) and low-SAA (N=20)). **(C)** Individual fold change of SSC levels in spot-urine of after control or low-SAA diet. Unpaired T-test of low-SAA compared to control arm (*\*P < 0.05*) was used in (A)-(C). All data are represented as mean ± standard deviation and each patient is depicted by a single dot. Abbreviations: h: hour; SAA: Sulfur-containing Amino Acids; spot-urine: spontaneous urine; SSC: S-sulfocysteine.

Conclusively, we could recapitulate adaptions in cysteine and sulfite metabolism in humans upon a low-SAA diet indicating conserved diet-induced mechanisms of stress-resistance.

## Discussion

Impaired cellular stress-resistance contributes to various diseases including common and life-threatening AKI, a frequent age-associated complication. Although CR effectively augments resilience leading to amelioration of AKI and superior survival rate in rodent models, a better understanding of the underlying mechanisms is required for successful clinical translation (8-11, 28, 38). Based on recent results regarding novel dietary regimens derived from longevity studies in animal models, five tailored dietary interventions were considered promising for the prevention of AKI beyond CR and their efficacy was systematically examined in a direct comparison (13, 14, 18, 20, 25, 39, 40). All five interventions had not been studied as protective strategies against AKI before and had never been directly compared to identify the most potent approach as well as associated metabolic patterns.

Regarding kidney function, survival, general health status and tissue damage, we identified FMD, and the low-SAA diets SR80/100 as the most potent preconditioning strategies beyond CR (32, 33). KD revealed less profound protective effects, but was indeed accompanied by improvements in survival and general health. In contrast, dietary mTOR inhibition by BCAA restriction failed to protect from renal ischemic damage and did not improve any of the indicators examined. Despite its beneficial effects on lifespan, inhibition of mTOR activity during the reperfusion in renal IRI using rapamycin has been shown to slow down kidney recovery after AKI, which may also explain the failure in renal protection by BCAA preconditioning we observed (41).

Considering that a reduction of SAA was among the most efficient approaches, we decided to examine the impact of the protective regimens on the TSP and cysteine metabolism. TSP enzyme-mediated generation of H_2_S is evolutionary conserved and plays a pivotal role in augmenting cellular stress-resistance and healthspan extension (1). The well-known protective effects of CR in ischemic hepatic damage, for example, are - at least in part - conveyed by the modulation of cysteine catabolism and production of H_2_S (23). The primary source of H_2_S generation are various metabolites that arise from cysteine metabolism. In addition to the H_2_S-producing branch, there are two further pathways involved in cysteine metabolism, (i) oxidative cysteine catabolism to taurine or sulfate and (ii) glutathione biosynthesis. Mechanistically, supplementation of the SAA methionine and cysteine abrogates the increased production of H_2_S and counteracts CR-mediated stress-resistance in rodents (23).

The most potent dietary regimens in our study - FMD, SR80/100 and CR - resulted in strong adaptations in oxidative and H_2_S-dependent cysteine catabolism. Despite the fact that the changes upon all of these four regimens point towards an overall increase in cysteine catabolism, the underlying mechanisms may differ as indicated by the distinct alterations of cysteine, homocysteine, sulfite, SSC and thiosulfate. All beneficial diets induce an elevation in serum thiosulfate with FMD showing the highest and most significant rise (>3fold). Importantly, elevated thiosulfate is a direct proxy for increased H_2_S turnover (23). This rise in thiosulfate after FMD is accompanied by a reduction in homocysteine and an elevation in cysteine, revealing a flux towards cysteine. Moreover, elevated levels of sulfite and thiosulfate indicate increased oxidative and H_2_S-dependent cysteine catabolism in response to its abundance after FMD. We could recapitulate the FMD-induced activation of oxidative cysteine catabolism on the protein level and show a strong upregulation of the initial and rate-limiting enzyme CDO1 - a prerequisite for the production of sulfite (Fig. 4). Furthermore, we identified an increase of the downstream enzymes GOT1/2 in response to FMD that directly accept cysteine and cysteine sulfinate for transamination contributing to H_2_S and sulfite generation, respectively, and are, consequently, both required for thiosulfate formation (Fig. 4, S8) (36). Similarly, both low-SAA diets revealed a rise in serum thiosulfate indicating the generation of H_2_S accompanied by an upregulation of MPST after SR100. As MPST desulfurizes 3-MP and transfers sulfane sulfur to its thiophilic acceptors, H_2_S is released. This rise in MPST was accompanied by a downregulation of key enzymes in cysteine catabolism outside the H_2_S-dependent branch, CSAD and GSS, leading to a shift in favor of the H_2_S-generating pathway (Fig. 4). However, in contrast to FMD and CR, SR100 resulted in a significant decrease of sulfite. While this may appear counterintuitive at first sight, our further metabolite quantifications provide a reasonable explanation for this finding. Of note, both low-SAA diets resulted in a substantial increase of urinary SSC, a secondary end product induced upon sulfite elevation. SSC is generated through the spontaneous, non-catalyzed reaction of sulfite with cystine and is widely used as a biomarker for sulfite levels (37, 42). High SSC values measured in SR80/100 indicate that the sulfite-producing H_2_S catabolism is indeed not only activated by FMD and CR but also by low-SAA diets. During the formation of SSC from sulfite and cystine, cysteine is released explaining the missing reduction (SR80) or even the rise in cysteine (SR100), even though dietary SAA supply was heavily reduced (Fig. S7) (43). Furthermore, this reaction also provides an explanation for the low sulfite levels observed in SR80/SR100 treated animals. This reciprocal relation of low sulfite and elevated SSC is not seen in CR and FMD diets.

Importantly, we confirmed diet-induced metabolic adaptations in cysteine metabolism in human bio-samples after SAA restriction. The rise in SSC generation was accompanied by a reduction in serum cystine and maintained cysteine levels upon dietary restriction of SAA in our pilot-cohort. This reduction in cystine is likely to be a consequence of its reaction with sulfite indicating active sulfite metabolism and cysteine maintenance as a shared pattern between rodents and humans in response to low-SAA diets. Sulfite is a common intermediate in oxidative and H_2_S-dependent cysteine catabolism and contributes to the reported H_2_S related signaling effects in cellular stress-resistance, i.e. by liberating H_2_S when reacting with persulfides (44–46). Additionally, sulfite itself is known to mediate cytoprotective effects by e.g., scavenging of reactive oxygen (ROS) and nitrogen species (RNS) or maintaining levels of cysteine and glutathione (1, 44, 45).

Currently, it remains unclear whether the benefits of an elevated cysteine catabolism including sulfite and H_2_S generation require a simultaneous increase in autophagy (23). In yeast, longevity benefits by methionine restriction are based on autophagy, whereas in mammals this remains unknown (47). Interestingly, we found strong hints using comparative proteome profiling, that FMD, SR80/100 and CR interfere with autophagy-associated genes in a similar fashion. Of note, three of the seven overlapping proteins (ATG7, PRKA and CTPS) that are differentially regulated by all of these four diets contribute to autophagy. Interestingly, conditional ATG7 ablation from proximal tubules aggravates AKI, whereas activation of autophagy protects from AKI (48, 49). The PKR (also named EIF2AK2) axis is activated by PRKA in the absence of double-stranded RNA (dsRNA), which is found during endoplasmic reticulum stress and in settings of insulin resistance (50). Conversely, low PRKA levels and a downregulated PKR pathway induce autophagy and protect from inflammation and oxidative stress (51). CTPS2, one of the two isoforms of the CTP synthase, is the key regulatory enzyme in pyrimidine biosynthesis. Interestingly, CTPS2 has an essential role in the maintenance of genome integrity and the synthesis of membranes (52).

Our study has several limitations: First, findings from our rodent IRI-model may not entirely reflect the pathogenesis of AKI in the clinical setting. Second, due to restraints regarding their production, SR80/100 and BCAA have an altered macronutrient ratio compared to the standard chow, which may additionally influence the given results. Third, as male rodents are more vulnerable to IRI we only examined male animals. Consequently, sexual dimorphism regarding dietary interventions is not reflected by our study. However, this aspect may also differ between mice and humans since the sex-related differences in susceptibility generally observed in mice are not obvious in available human data (39, 53). Fourth, direct pharmacological manipulation of sulfite and H_2_S would be an attractive target for clinical translation and will need to be addressed in follow-up studies (54).

Regarding our human data, the limited number of participants in our pilot-cohort recapitulating the observed metabolic adaptions of the beneficial diets has to be acknowledged. Furthermore, the dietary intervention was executed in an outpatient setting associated with potential issues regarding adherence. Nonetheless, the key finding regarding SSC was indeed confirmed in a larger cohort.

In conclusion, our systematic and direct comparison of five tailored dietary preconditioning regimens identified FMD and SAA-restriction as highly potent strategies for organ protection beyond CR. We identified modulation of cysteine catabolism as a pivotal common aspect of these interventions with sulfite metabolism as its central component. The impact of SAA-restriction on cysteine metabolism could be confirmed in a human cohort. Since both FMD and low-SAA diets are feasible in humans our findings provide an important outlook towards novel protective strategies in the patient setting. Besides dietary interventions, pharmacological modulation of oxidative and H_2_S-dependent cysteine catabolism may be exploited in the future.

## Methods

### Dietary Preconditioning in Rodents

Male C57BL6N mice were obtained at 6 weeks of age from Janvier Labs (Paris, France) and maintained individually upon arrival in an identical specific pathogen free environment with constant temperature and humidity and 12h/12h light/dark cycle. Unless otherwise on specific experimental diets, mice were fed *ad libitum* with their regular chow (V155-330 ssniff GmbH, Soest, Germany) and had unrestricted access to water.

Male C57BL6N mice aged 8 weeks were allocated randomly to their specific dietary regimen and transferred to a fresh cage at beginning of dietary preconditioning. FMD, SR80/100, KD, BCAA and CR were performed for 14 days prior to renal IRI simultaneously (Fig. 1A). Rodent periodic fasting with FMD is a 3-day regimen and was performed as previously described (13, 14). Day 1 of FMD (S5159-E736, ssniff GmbH, Soest, Germany) consists of a combination of flavored broth mixes, vegetable powders, extra virgin olive oil (EVOO), essential fatty acids, minerals and vitamins, whereas on day 2-3 of FMD a combination of flavored broth mixes, glycerol and hydrogel were supplied to the mice (S5159-E738, ssniff GmbH, Soest, Germany) (Table S4). On day 4-7 of FMD mice had *ad libitum* access to their standard chow. Energy intake was reduced to 7.67 kJ/g on day 1 and 1.48 kJ/g on day 2-3 of FMD, respectively. Rodent FMD regimen was performed twice before renal IRI and rodent surgery was performed after the second period with an *ad libitum* access to the standard chow on day 14. KD (S51519-E730, ssniff GmbH, Soest, Germany) contained 10% protein, 89% fat and 0.6% carbohydrates and mice were supplied with KD daily and had *ad libitu*m access to the high-fat chow (Table S5). BCAA chow (S5159-E724, ssniff GmbH, Soest, Germany) resulted in a 66% reduction of leucine, isoleucine and valine as previously described (20). To reduce the daily uptake of SAA, either a full restriction of methionine and cysteine (SR100, S5159-E720, ssniff GmbH, Soest, Germany) or an 80%-reduction of SAA (SR80, S5159-E722, ssniff GmbH, Soest, Germany) were used (Table S5). BCAA and the low-SAA diets SR80/100 were supplied weekly and mice had *ad libitum* access to the respective chow. Animals preconditioned by CR received 70% of the regular daily intake (3g/d) of their regular chow for 14 days prior to renal IRI and they were supplied with food in the morning hours (9.00 - 11.00 am) on each day during dietary regimen. Food intake for CR animals was chosen based on previous rodent studies examining CR in the context of AKI (9, 10). CR animals were not fed on the day of surgery.

During dietary preconditioning mice had *ad libitum* access to water and they consumed all supplied diets revealing no signs of food aversion. To assess weight loss mice were weighted weekly and no increase in morbidity or mortality was observed during any of the dietary interventions.

### Renal Ischemia-Reperfusion Injury Model

To induce AKI, a warm renal ischemia-reperfusion ischemia (IRI) model as previously described with slight modifications was used (9). Briefly, to perform surgery, mice were anesthetized with i.p. administered ketamine (Zoetis Inc. Parsipanny, NJ, USA) / xylazine (Bayer AG, Leverkusen, Germany). A thermometer was inserted rectally and mice were placed on a temperature-controlled heating pad (Havard apparatus, Holliston, MA, USA) to maintain body temperature. After midline laparotomy the right kidney was removed and the left renal vessels were clamped for 40 minutes with an atraumatic micro-vascular clamp. Renal ischemia was verified visually by color change of the kidney and afterwards the abdomen was covered with a compress soaked in 0.9% saline solution. After restoration of renal blood flow was inspected visually, the abdomen was closed in two layers and the mice were placed in a fresh cage. For analgesia the mice received buprenorphine (Indivior Europe Limited, Dublin, Ireland), in a dosage of 0.1mg/kg per body weight injected s.c. twice daily, as well as 0.009mg/mL buprenorphine administered in their drinking water. After surgery, dietary preconditioned and non-PC mice had *ad libitum* access to the standard chow.

CR served as positive control, non-PC animals as negative. Another experimental control consisted of sham animals, which underwent right nephrectomy without clamping of the vascular pedicle of the contralateral kidney.

Mice were euthanized after 24 hours or 72 hours after reperfusion. Therefore, mice were anesthetized with i.p. ketamine / xylazine injection and final blood withdrawal was performed by puncture of the right ventricle. Intraoperative and postoperative obtained kidneys were cut into two equal sections, which were either snap frozen in liquid nitrogen or fixed in 4% formaldehyde and subsequently embedded in paraffin, respectively. Baseline animals were sacrificed by final blood drawing after dietary interventions but before renal IRI. Both kidneys were cut into two equal sections and these were either snap frozen in liquid nitrogen or fixed in 4% formaldehyde with subsequent paraffin embedding. Mouse urine was directly snap-frozen in liquid nitrogen upon collection. Mouse urine and serum samples were stored at −80°C.

### Renal Function and Health Performance after Renal Ischemia-Reperfusion Injury

Mouse serum creatinine and urea levels were measured by the central laboratory of the University Hospital Cologne using a Cobas C 702 (Roche Diagnostics GmbH, Penzberg and Mannheim, Germany). Postoperative health performance was examined in the morning hours (9.00 - 11.00 am) of each day using a governmental authorities approved recovery score that consisted of weight loss, appearance and activity (Table S1). Mortality and postoperative recovery were assessed over a period of 72 hours after renal IRI.

### Histopathology

Formalin-fixed, paraffin-embedded tissue was cut into 2-μm-thick slices and stained with periodic acid–Schiff (PAS). Acute tubular damage was classified in a semi-quantitative and blinded manner by an experienced nephropathologist (H.G.) using a composite score assessing vacuolization, epithelial flattening, loss of either brush borders or nuclei and tubular necrosis (32, 33). According to the affected area grades were given from 0-4 (1: 0%–25%, 2: 25%–50%, 3: 50%–75%, 4: 75%–100%). Images of PAS-staining were obtained using a slide scanner SCN4000 (Leica, Wetzlar, Germany) in a 20× magnification.

### Terminal Deoxynucleotidyl Transferase–Mediated Digoxigenin-Deoxyuridine Nick-End Labeling Staining

DeadEnd Fluorometric TUNEL System (Promega, Fitchburg, WI, USA) was used on 2 µm thick formalin-fixed paraffin sections according to the manufacturer’s protocol. Images were taken on an Axiovert 200M microscope with a Plan-Apochromat 20×/0.8 objective and an AxioCamMR camera (all Carl Zeiss MircoImaging GmbH, Jena, Germany).

### Immunohistochemistry

Cleaved-caspase-3 immunostaining was performed on 2 µm thick formalin-fixed paraffin sections in a 1:200 dilution according to the manufacture’s protocol (Cell Signaling Technology, Danvers, MA, USA). Pictures were taken on a slide scanner SCN4000 in a 20× magnification.

### Human samples

Human biosamples were collected from patients participating in a randomized placebo-controlled trial after 7 days of dietary preconditioning using formula diets; trial design is described in detail on clinicaltrials.gov (NCT03715868). Briefly, the control arm was fed a standard liquid formula diet, whilst the intervention arm received a liquid formula diet containing SAA levels reduced by 77%. Participants were randomly assigned to the two treatment groups. Daily caloric intake was not restricted and equal in both groups.

Urine creatinine concentrations were measured in the central laboratory of the University Hospital Cologne using a Cobas C 702. All urine-samples for further biochemical analysis were stored at −80°C. Plasma cystine was determined in the central laboratory of the University Hospital Cologne in pre-dietary, as well as post-dietary samples obtained on the seventh day of dietary intervention. Metabolic characterization of cysteine catabolism was performed in a pilot-cohort of ten sex-and age-matched participants. Urinary SSC was additionally quantified as a read-out for cysteine catabolism in a larger cohort of 49 patients adhering to either a low-SAA or control diet.

### Targeted High-Performance Liquid Chromatography

Cysteine, homocysteine, sulfite, thiosulfate and s-sulfocysteine (SSC) were determined by targeted high-performance liquid chromatography (HPLC). For the detection of cysteine, homocysteine, sulfite and thiosulfate in murine and human serum or urine we chose a monobromobimane (mBBr) derivatization approach. Briefly, defrosted samples were centrifuged and the supernatant were mixed with 4-(2-hydroxyethyl)-1-piperazineethanesulfonic acid (HEPES), ethylenediaminetetraacetic acid (EDTA) and mBBr in acetonitrile. The reaction was performed in the dark and was stopped with methanesulfonic acid (MsOH). Before injection, the samples were diluted with four volumes of the respective 0.5% acetic acid used as running buffer A, and transferred into a glass HPLC vail. The samples were then transferred on a 125 x 4.0 mm phase 60RP-Select B LiChroCART 250-4 (Sigma-Aldrich St. Louis, MO, USA) and acetic acid was used as running buffer A, methanol as running buffer B. The gradient was: 0–4 min, 5% B; 4–8 min, 20% B; 8–15 min, 25% B; 15–27 min, 60% B; 27–28 min, 100% B; 28–32 min, 100% B; 32–33 min, 5% B; 33–60 min, 5% B. Dabsylated products were determined spectrophotometrically.

SSC was detected using a derivatization reaction with 9-fluorenylmethyloxycarbonyl chloride (Fmoc-Cl) as previously described (55). In brief, murine and human urine were centrifuged and the supernatant was mixed with sodium carbonate / bicarbonate (Na2CO3 / NaHCO3) and Fmoc-Cl in acetonitrile. The derivatization mixture was incubated in the dark and the reaction was stopped through the addition of amantadine-hydrogen chloride (HCl). All samples were centrifuged and the supernatant was transferred into a glass HPLC vial. SSC-Fmoc was separated on a Nucleodur HTec C18 RP column (Macherey-Nagel GmbH, Düren, Germany). Running buffer A was Na-formate, trifluoroacetic acid and running buffer B was methanol. SSC was eluted at 1 mL/min in buffer B at 35-65% and 0-15.0 min gradient, eluting at 9.97 min. SSC was quantified based on area under the peak at 265 nm.

### Absorption-based creatinine determination in mouse urine

Thawed murine urine samples were diluted 1:10 in HCl and transferred together with standard curve of creatinine into individual wells of a 96-well plate. After incubating with alkaline picrate solution in the dark, urine creatinine concentration were determined spectrophotometrically.

### Proteomic analysis with Liquid Chromatography and Tandem Mass Spectrometry

All samples were analyzed on a Q Exactive Plus Orbitrap mass spectrometer coupled to an EASY nLC 1000 (both Thermo Fisher Scientific, Waltham, MA, USA). Peptides were loaded with solvent A (0.1% formic acid in water) onto an in-house packed analytical column filled with Poroshell EC120 C18 material (Agilent Technologies, Santa Clara, CA, USA). Peptides were chromatographically separated at a constant flow rate of 250 nL/min using the following gradient: 3-4% solvent B (0.1% formic acid in 80 % acetonitrile) within 1.0 min, 4-27% solvent B within 119.0 min, 27-50% solvent B within 19.0 min, 50-95% solvent B within 1.0 min, followed by washing and column equilibration. The mass spectrometer was operated in data-dependent acquisition mode. The MS1 survey scan was acquired from 300-1750 m/z at a resolution of 70,000. The top 10 most abundant peptides were isolated within a 1.8 Th window and subjected to HCD fragmentation at a normalized collision energy of 27%. The AGC target was set to 5e5 charges, allowing a maximum injection time of 55 ms. Product ions were detected in the Orbitrap at a resolution of 17,500. Precursors were dynamically excluded for 30.0 s. Raw data were processed with MaxQuant (version 1.5.3.8) using default parameters searching against the canonical *Mus musculus* reference proteome (UP000000589) downloaded on the 26th of August, 2020. Results were further analyzed in Perseus 1.6.1.1 and RStudio (RStudio Inc., Boston, MA, USA) as previously described (10). Briefly, proteins identified by only one site and reverse hits were removed. Label-free quantification of proteins were log_2_ transformed and filtered for at least 4 valid measured values being present per dietary group. Two-tailed *t*-tests and two-tailed analysis of variances (ANOVA) were calculated where appropriate. To determine significant proteins and to correct for multiple testing were a FDR was set to 0.05. Functional data were plotted using RStudio and the ggplot2 as well as cowplot packages.

### Statistics

Statistical analysis was done with GraphPad Prism software version 5 (GraphPad Prism Software Inc. San Diego, CA, USA). Animals were randomly allocated to their dietary preconditioning group. Rodent experiments were not blinded due to the different diets. The histological damage score, biochemistry analyses as well as serum creatinine and serum urea measurements were conducted blinded. Samples sizes were chosen based on experience with previous studies employing renal IRI in rodents (n=10 and n=13 mice per time-point and dietary regimen, respectively) and no power analysis was done to determine samples sizes (8, 9). For metabolic and proteome profiling at least four biologic replicates were tested. Ten adults that underwent dietary SAA restriction using a formula diet for seven days compared to ten age- and sex-matched individuals on a formula diet containing standard levels of SAA were used as a pilot-cohort to confirm metabolic adaptions in humans. Human participants were randomly assigned to the two treatment groups in a double-blinded clinical trial. In four adults, spot- and 24h-urine collection of either pre- or post-diet was missing. Urinary SSC was additionally quantified in a larger cohort of 49 patients. No data nor outliers were excluded from the study. Results are depicted as mean ± standard deviation. One-way ANOVA and Bonferroni multiple comparison test were used to examine differences between groups unless otherwise indicated. *P < 0.05* was considered significant.

### Study approval

All animal procedures were approved by the State Agency for Nature, Environment and Consumer Protection North Rhine-Westphalia, Germany, (TVA2019.A345). The human study protocol was approved by the Institutional Review Board of the University Hospital Cologne and informed consent was obtained from all participants prior to inclusion. Trial design is described in detail on clinicaltrials.gov (NCT03715868).

## Author contributions

F.C.K. and C.Y.F. performed experiments, F.C.K., C.Y.F., M.R.S., K.J.R.H.-A., K.B., H.G., J.S., A.A., G.S., V.B. and R.-U.M. analyzed experiments, F.G., T.O., C.G., R.-U.M. and V.B. contributed human bio-samples, F.C.K., V.B. and R.U.M. designed the study, J.-W.L. and G.S. contributed new tools, F.C.K., G.S, V.B. and R.-U.M interpreted results, F.C.K, G.S., V.B. and R.-U.M. wrote the article. M.R.S., K.J.R.H.-A., F.G., J.S., A.A., T.B., and B.S. revised the manuscript.

## Acknowledgments

FCK reports grants by Else Kröner-Fresenius-Stiftung, by the German Research Foundation under Germany’s Excellence Strategy - EXC 2030: CECAD - Excellent in Aging Research - Project number 390661388 and by the Koeln Fortune program / Faculty of Medicine, University of Cologne during the conduct of this project. MRS receives funding from the Koeln Fortune program and by the Cologne Clinician Scientist Program (CCSP) / Faculty of Medicine / University of Cologne. Funded by the German Research Foundation (DFG, FI 773/15-1)./ Faculty of Medicine, University of Cologne. FG is supported by the Koeln Fortune program / Faculty of Medicine, University of Cologne and the German Research Foundation (GR3932/2-1). R.-U.M. was supported through the Nachwuchsgruppen.NRW program of the Ministry of Science North Rhine-Westphalia and received further funding from the German Research Foundation (CRU 329, MU 3629/3-1).

We thank Serena Greco-Torres for her excellent technical assistance. Special thanks to the *in-vivo* Research Facility at the Cologne Excellence Cluster on Cellular Stress Responses in Aging-Associated Diseases (CECAD) for help with the mouse studies. Furthermore, we acknowledge the support by the study-center of the Department II of Internal Medicine, University Hospital of Cologne. The authors dedicate this work to James R. Mitchell for his ground-breaking research in dietary restriction and hydrogen sulfide metabolism, who tragically passed away last year.

