## Supplemental Material for "A systematic analysis of diet-induced nephroprotection reveals overlapping and conserved changes in cysteine catabolism"

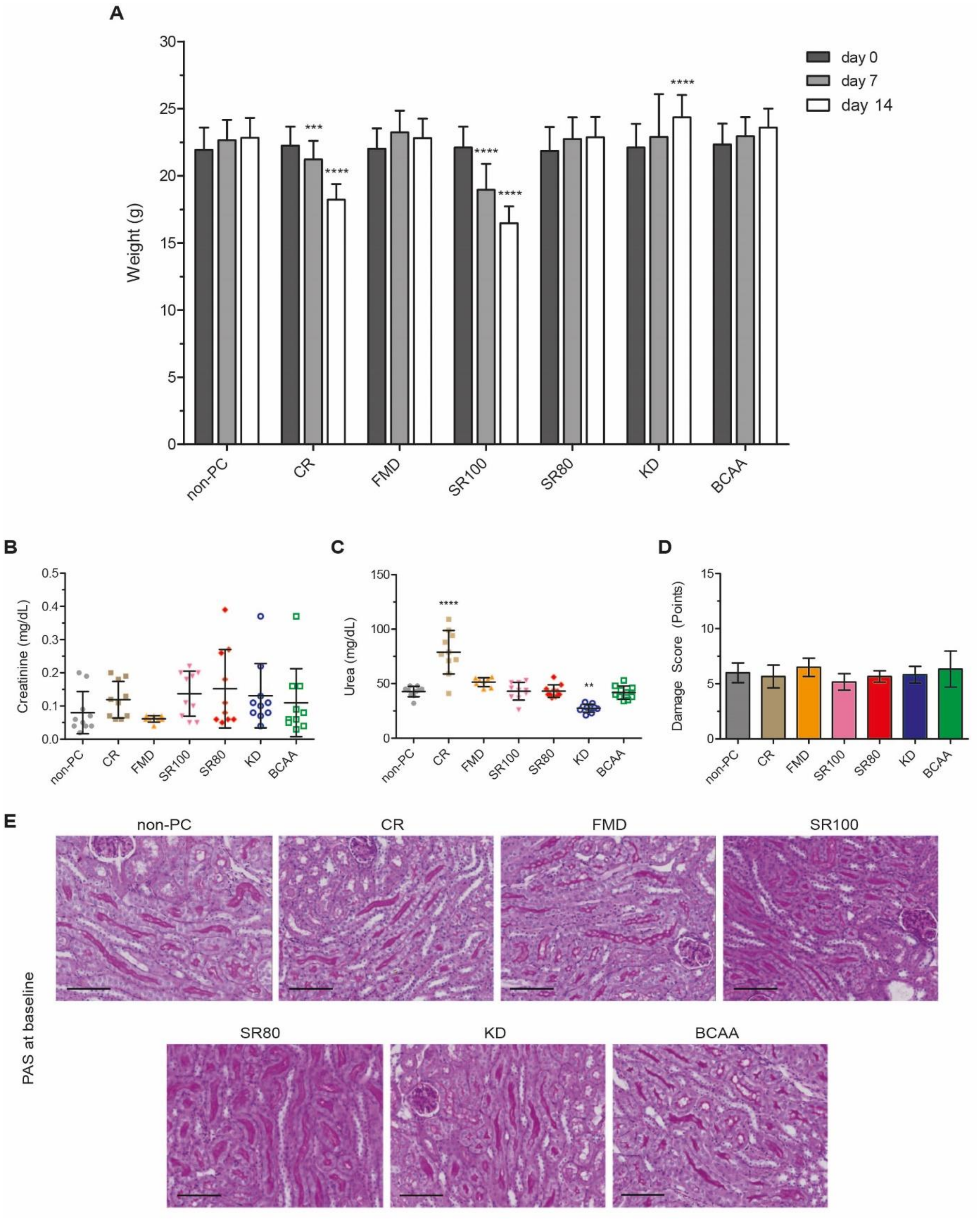


**Fig. S1. Phenotyping and functional kidney parameters after tailored dietary preconditioning before renal IRI.**

**(A)** Weight development during dietary preconditioning before IRI. **(B, C)** Creatinine and urea concentrations for dietary preconditioned and non-PC animals at 0 hours after dietary preconditioning before renal IRI. **(D)** Histologic damage score at 0 hours after dietary preconditioning before renal IRI. One-way ANOVA and Bonferroni multiple comparison test across all dietary preconditioning groups indicating differences to non-PC mice (***P < 0.01, ***P < 0.001* and *****P < 0.0001*) was used in (A)-(D). Data are represented as mean ± standard deviation and n = 10 per dietary group. **(E)** Kidney histology by PAS-stainings in dietary preconditioned animals at 0 hours after dietary preconditioning before renal IRI (. Abbreviations: BCAA: Branched Chain Amino Acids; CR: Caloric Restriction; FMD: Fasting Mimicking Diet; KD: Ketogenic Diet; non-PC: non-preconditioned animals; PAS: Periodic acid-Schiff; SR100: 100% Restriction of Sulfur-containing Amino Acids; SR80: 80% Reduction of Sulfur-containing Amino Acids.

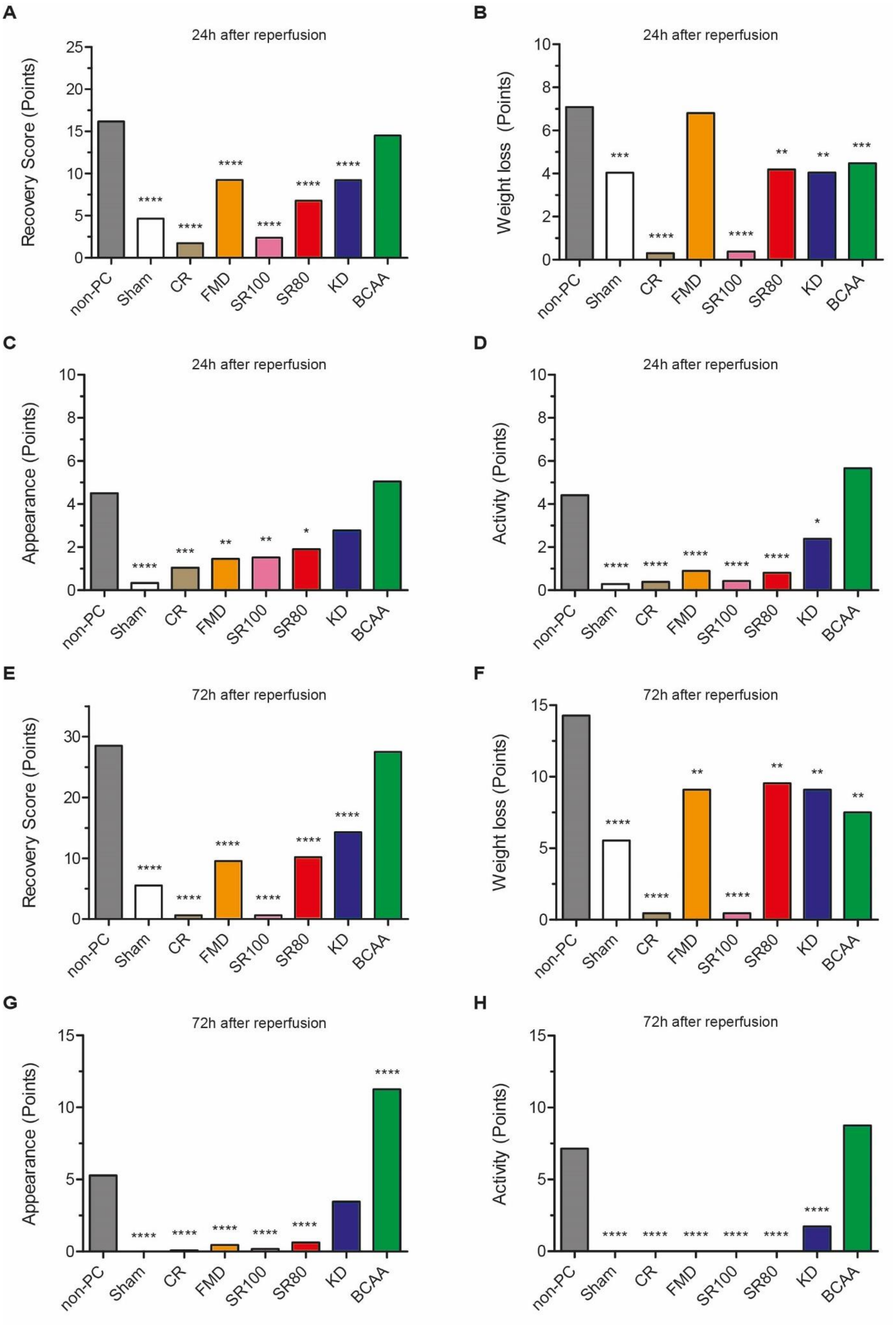
A
24h after reperfusion
Recovery Score (Points)
B
24h after reperfusion
Weight loss (Points)
C
24h after reperfusion
Appearance (Points)
D
24h after reperfusion
Activity (Points)
E
72h after reperfusion
Recovery Score (Points)
F
72h after reperfusion
Weight loss (Points)
G
72h after reperfusion
Appearance (Points)
H
72h after reperfusion
Activity (Points)
non-PC
Sham
CR
FMD
SR100
SR80
KD
BCAA

**Fig. S2. FMD, SR80/100, KD and CR lead to better recovery after renal IRI.**

The recovery score after renal IRI consisted of the three categories weight loss, activity and appearance (Table S1). Recovery score and its respective sub-items weight, appearance and activity examined **(A-D)** 24 hours, as well as **(E-H)** 72 hours after reperfusion, respectively. Lower scores indicate better recovery. One-way ANOVA and Bonferroni multiple comparison test across all dietary preconditioning groups indicating differences to non-PC mice (*$P < 0.05$, **$P < 0.01$, ***$P < 0.001$ and ****$P < 0.0001$). All data are represented as means. 24h after IRI: n = 10 per dietary group; 72h after IRI n = 13 in the FMD, SR80/100, CR and sham groups, n = 9 in the KD, n = 4 in the BCAA and n = 6 in the non-PC group, respectively. Abbreviations: BCAA: Branched Chain Amino Acids; CR: Caloric Restriction; FMD: Fasting Mimicking Diet; KD: Ketogenic Diet; non-PC: non-preconditioned animals; SR100: 100% Restriction of Sulfur-containing Amino Acids; SR80: 80% Reduction of Sulfur-containing Amino Acids.

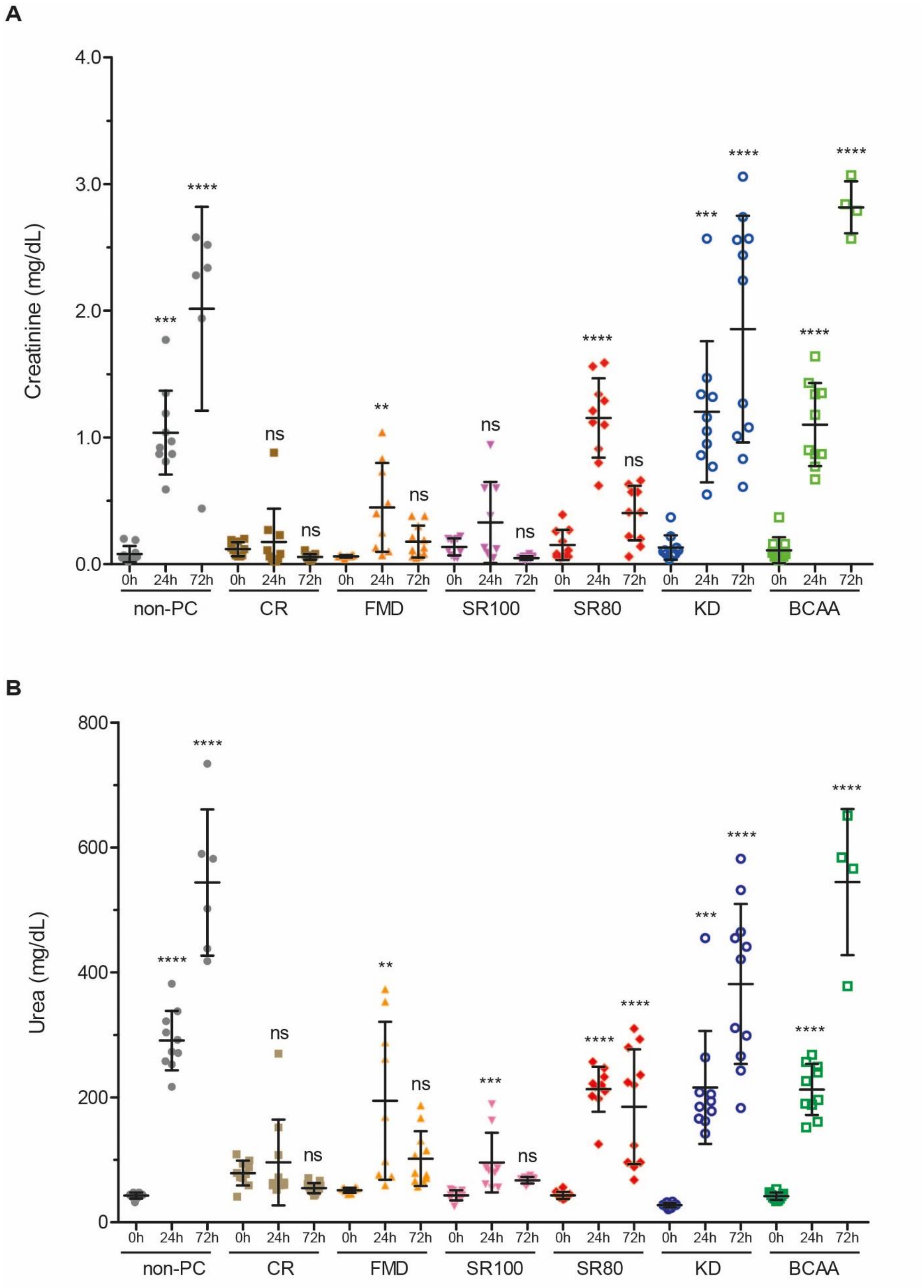


**Fig. S3. Kinetics in creatinine and urea concentrations after renal IRI revealing preconditioning mediated protection against AKI induced by FMD, SR80/100 and CR.**

**(A)** Creatinine concentrations of dietary preconditioned and non-PC animals at baseline, 24 hours and 72 hours after reperfusion**(B)** Urea concentrations of dietary preconditioned and non-PC animals at baseline, after 24 hours and 72 hours after reperfusionOne-way ANOVA and Bonferroni multiple comparison test across all dietary preconditioning groups indicating differences to baseline animals of the respective treatment group . All data are represented as mean ± standard deviation and each mice is depicted by a single dot. Abbreviations: BCAA: Dietary Restriction of Branched Chain Amino Acids; CR: Caloric Restriction; FMD: Fasting Mimicking Diet; h: hour; KD: Ketogenic Diet; non-PC: non-preconditioned animals; SR100: 100% Restriction of Sulfur-containing Amino Acids; SR80: 80% Reduction of Sulfur-containing Amino Acids.

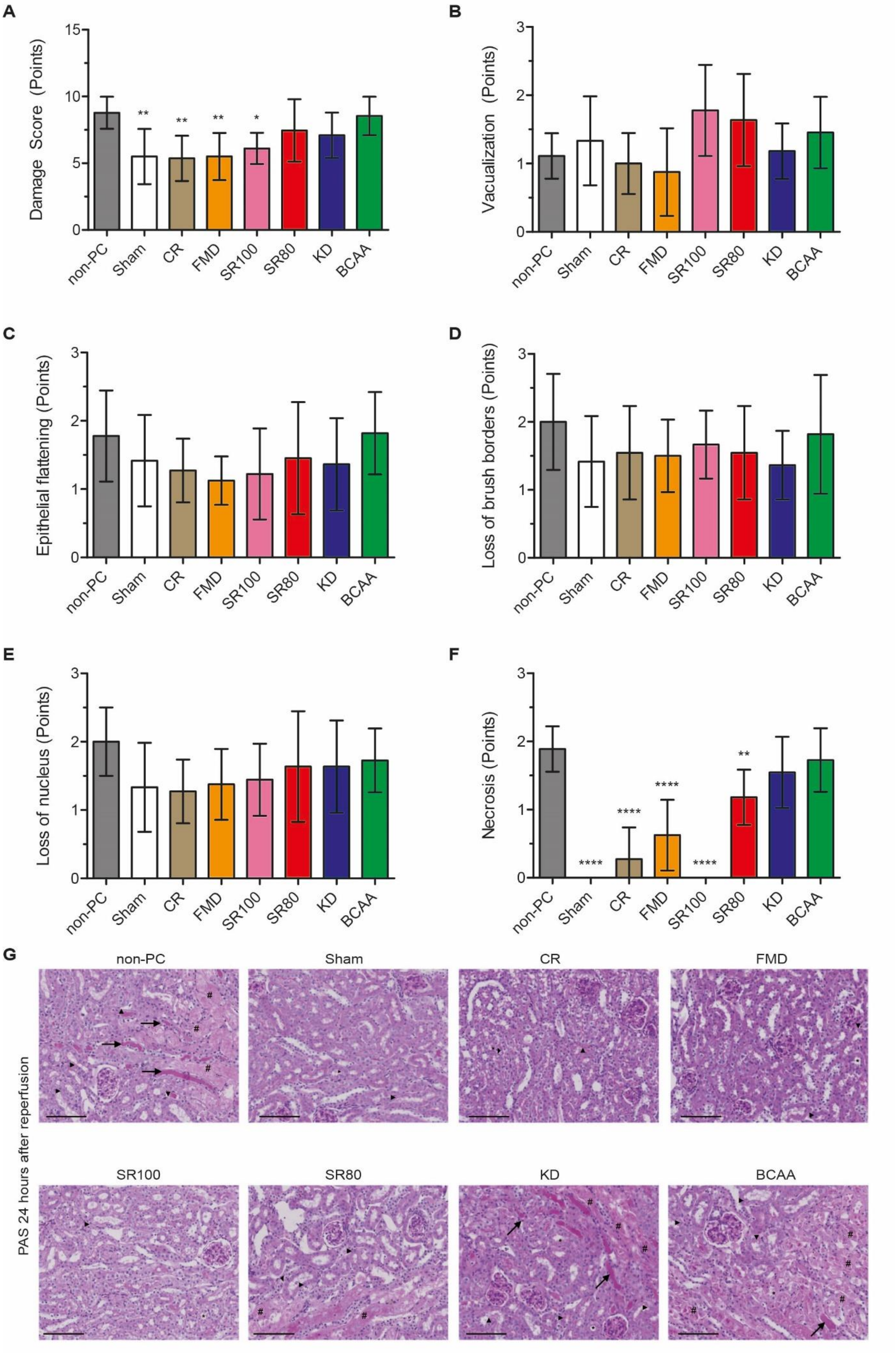
A
Damage Score (Points)
B
Vacualization (Points)
C
Epithelial flattening (Points)
D
Loss of brush borders (Points)
E
Loss of nucleus (Points)
F
Necrosis (Points)
non-PC
Sham
CR
FMD
SR100
SR80
KD
BCAA
G
PAS 24 hours after reperfusion

**Fig. S4. FMD, SR80/100 and CR protect form tubular necrosis quantified by histopathology after renal IRI.**

**(A)** Histologic damage score and its respective subtitems **(B)** vacuolization, **(C)** epithelial flattening, **(D)** loss of brush borders **(E)** loss of nucleus and **(F)** tubular necrosis quantified at 24 hours after reperfusion in dietary preconditioned, non-PC and sham animals. One-way ANOVA and Bonferroni multiple comparison test across all dietary preconditioning groups indicating significant differences compared to non-PC ($*P < 0.05$, $**P < 0.01$ and $****P < 0.0001$) was used in (A)-(F). Data are represented as mean ± standard deviation and n = 10 per dietary group. **(** hours after reperfusion . Arrowheads indicate nucleolar loss, arrows tubular casts, * epithelial flattening and # denuded tubuli with luminar debris, respectively. Abbreviations: BCAA: Branched Chain Amino Acids; CR: Caloric Restriction; FMD: Fasting Mimicking Diet; KD: Ketogenic Diet; non-PC: non-preconditioned animals; PAS: Periodic acid-Schiff; SR100: 100% Restriction of Sulfur-containing Amino Acids; SR80: 80% Reduction of Sulfur-containing Amino Acids.

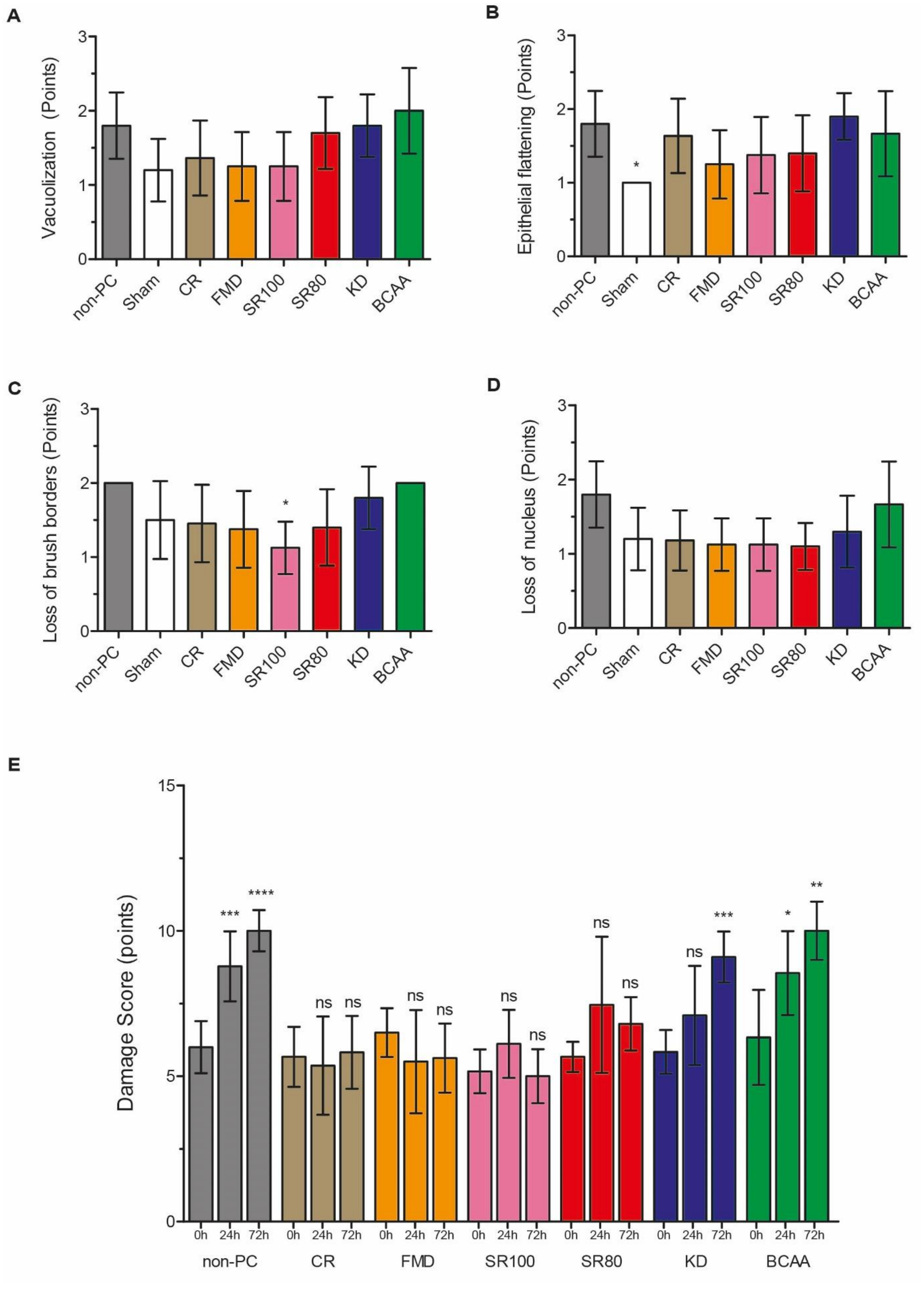


**Fig. S5. Subitems of a histologic damage score at 72 hours after reperfusion and longitudinal analysis of damages in kidney architecture after renal IRI.**

**(A)** Vacuolization, **(B)** epithelial flattening, **(C)** loss of brush borders and **(D)** loss of nucleus were quantified by experienced nephropathologist in a blinded manner in dietary preconditioned, non-PC and sham animals 72 hours after reperfusion. One-way ANOVA and Bonferroni multiple comparison test across all dietary preconditioning groups indicating significant differences compared to non-PC animals (*$P < 0.05$) was used in (A)-(D). Data are represented as mean ± standard deviation in (A)-(D). n = 13 in the FMD, SR80/100, CR and sham group, n = 9 in the KD, n = 4 in the BCAA and n = 6 in the non-PC group, respectively. **(E)** Longitudinal analysis of damages in kidney architecture after renal IRI as quantified by a composite histologic damage score in dietary preconditioned and non-PC animals at 0 hours, 24 hours and 72 hours after reperfusionOne-way ANOVA and Bonferroni multiple comparison test across all dietary preconditioning groups indicating differences to baseline animals of the respective treatment group. Data are represented as mean ± standard deviation. 0h (before IRI): n =10 per dietary group; 24h after IRI: n = 10 per group; 72h after IRI n = 13 in the FMD, SR80/100, CR and sham groups, n = 9 in the KD, n = 4 in the BCAA and n = 6 in the non-PC group, respectively. Abbreviations: BCAA: Branched Chain Amino Acids; CR: Caloric Restriction; FMD: Fasting Mimicking Diet; KD: Ketogenic Diet; non-PC: non-preconditioned animals; SR100: 100% Depletion of Sulfur Containing Amino Acids; SR80: 80% Reduction of Sulfur Containing Amino Acids.

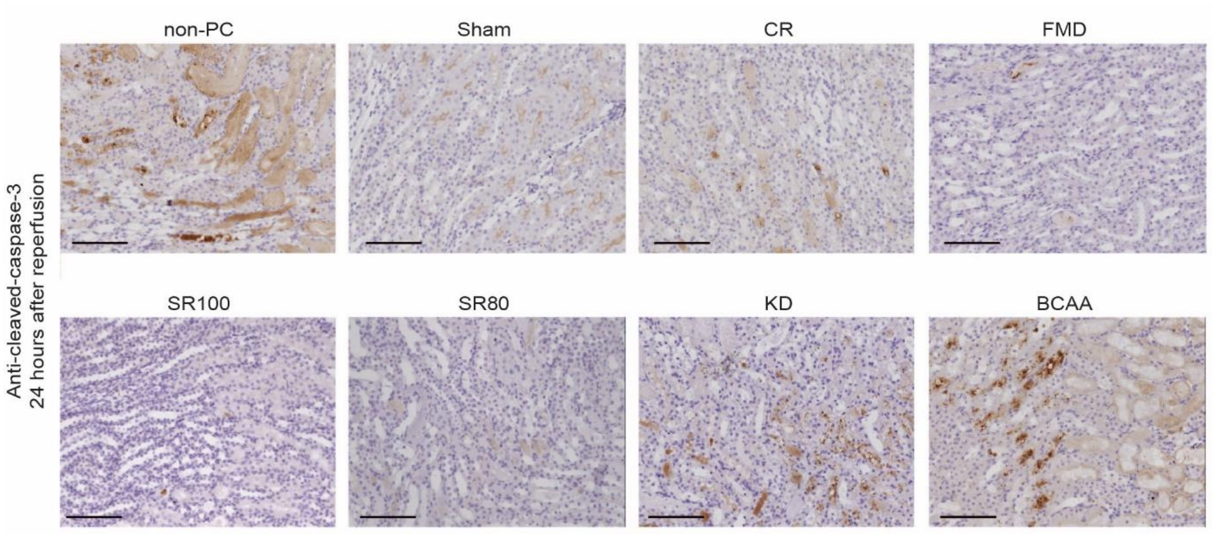


**Fig. S6. FMD, SR80/100 and CR reduce immunohistochemical signs of cell damage after renal IRI**.

Abbreviations: BCAA: Branched Chain Amino Acids; CR: Caloric Restriction; FMD: Fasting Mimicking Diet; KD: Ketogenic Diet, non-PC: non-preconditioned animals; SR100: 100% Depletion of Sulfur-containing Amino Acids; SR80: 80% Reduction of Sulfur-containing Amino Acids.

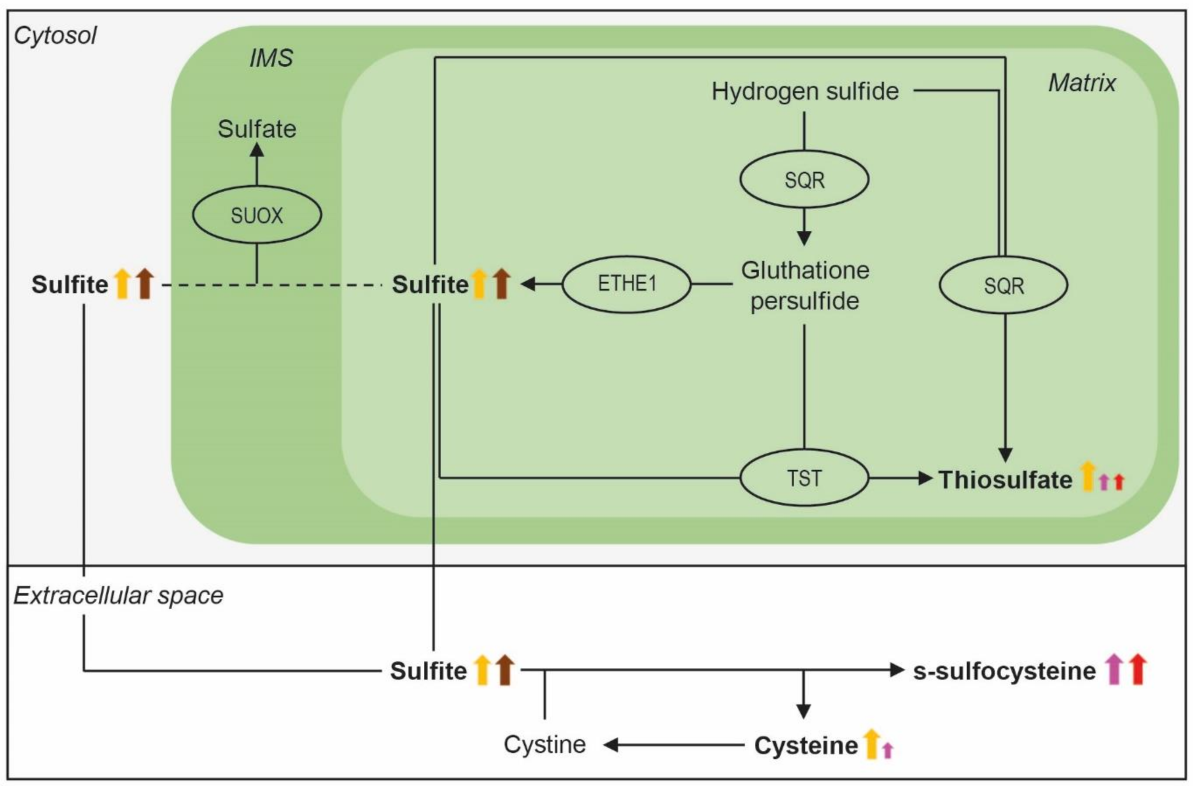


**Fig. S7. Schematic illustration of oxidative and hydrogen sulfide ($H_2S$)-dependent cysteine catabolism in response to FMD, SR80/100 and CR outlines a common central role for sulfite.**

Depicted are significant elevations of the key intermediate sulfite, the end-products thiosulfate and s-sulfocysteine as well as cysteine in response to FMD, SR80/100 and CR. Arrow's heights correspond to the quantity in change compared to non-preconditioned mice determined by high performance liquid chromatography. S-sulfocysteine (SSC) is generated by the non-catalyzed reaction of cystine and sulfite, which results in the additional formation of cysteine. SSC is a biomarker correlating with sulfite levels. Arrows' color indicate the dietary group (Fasting Mimicking Diet (FMD): orange; 100% Restriction of Sulfur-containing Amino Acids (SR100): pink; 80% Restriction of Sulfur-containing Amino Acids (SR80): red; Caloric Restriction (CR): brown). Enzymes are depicted as ellipsis. Arrows with solid line indicate enzymatic, arrows with dashed lines non-enzymatic reactions, respectively. The investigated major end-products of the $H_2S$-generating branch SSC and thiosulfate as well as the central intermediate sulfite are written in bold. Abbreviation: IMS: mitochondrial intermembrane space.

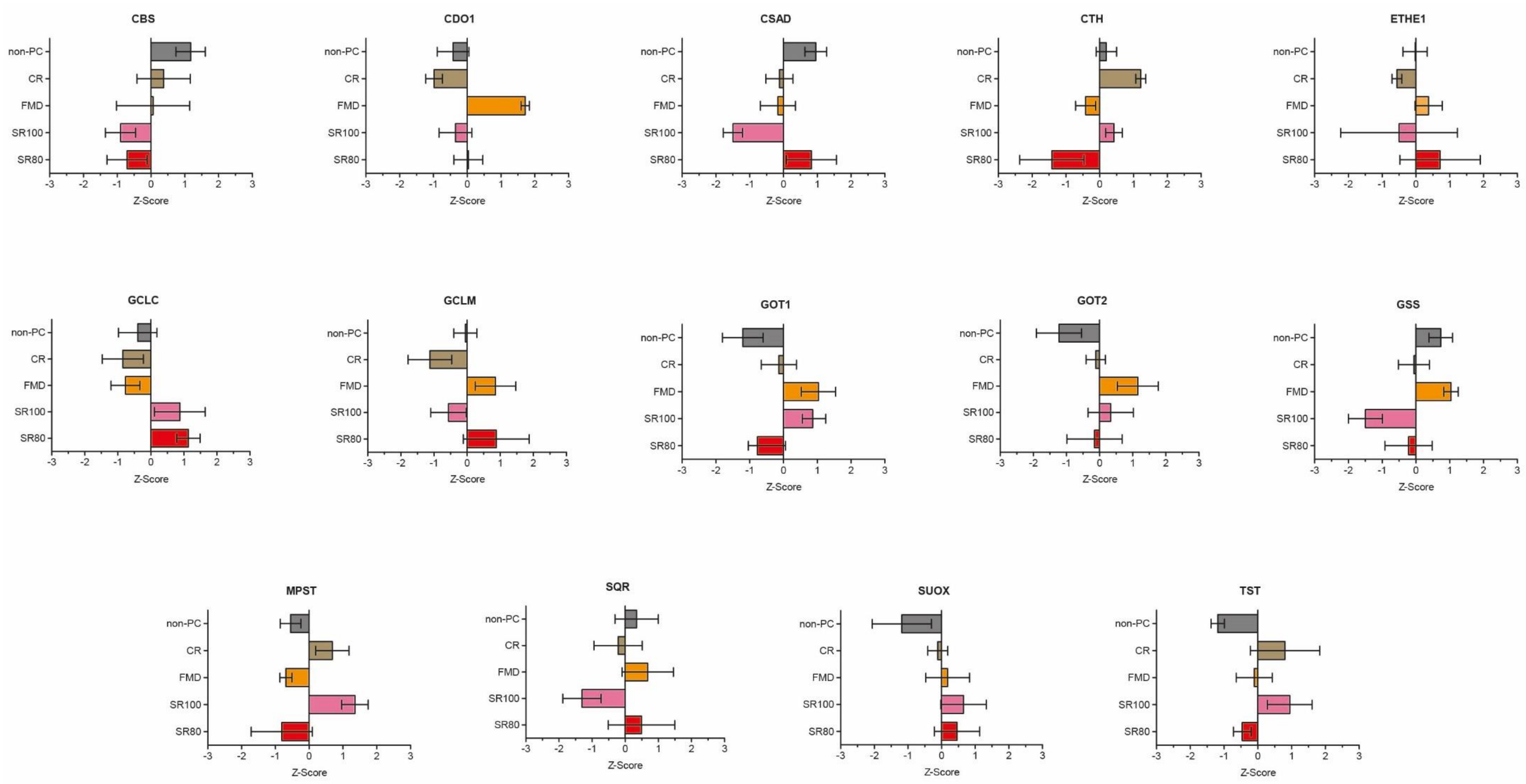
CBS
CDO1
CSAD
CTH
ETHE1
GCLC
GCLM
GOT1
GOT2
GSS
MPST
SQR
SUOX
TST
non-PC
CR
FMD
SR100
SR80
-3 -2 -1 0 1 2 3
Z-Score

**Fig S8. Specific renal enzymatic changes in oxidative and hydrogen sulfide ($H_2S$)-dependent cysteine catabolism in response to FMD, SR80/100 or CR quantified by comparative proteome profiling.**

Depicted are z-scores of key enzymes of oxidative and hydrogen sulfide ($H_2S$)-dependent cysteine metabolism. Z-scores of four biological replicates per dietary group are represented as mean ± standard deviation. Abbreviations: CR: Caloric Restriction; FMD: Fasting Mimicking Diet; non-PC: non-preconditioned animals; SR100: 100% Restriction of Sulfur-containing Amino Acids; SR80: 80% Reduction of Sulfur-containing Amino Acids.

A

B

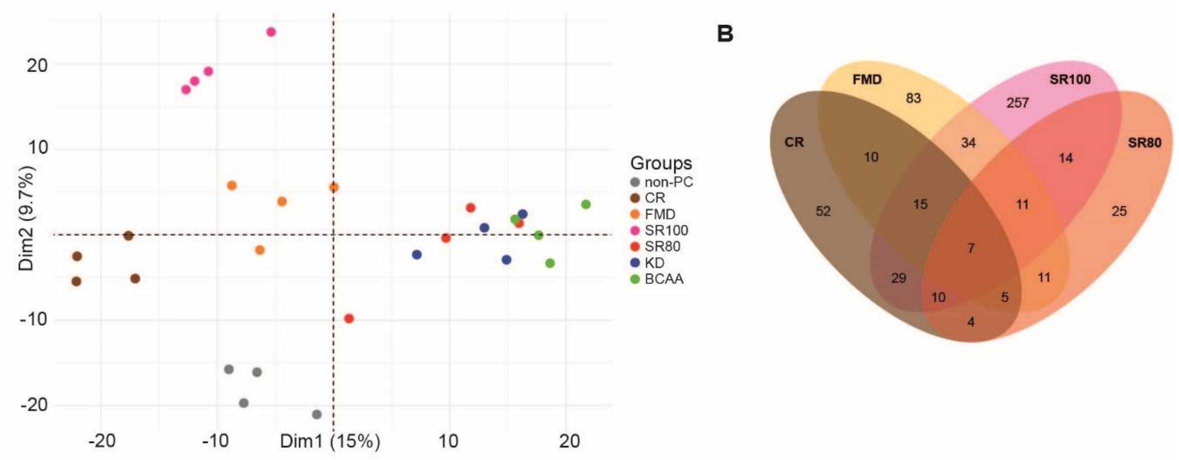


C

| | FMD | | SR100 | | SR80 | | CR | |
|---|---|---|---|---|---|---|---|---|
| | Log2FC | q-value | Log2FC | q-value | Log2FC | q-value | Log2FC | q-value |
| ATG7 | -1.87199 | 0.0251273 | -2.19311 | 0.0068108 | -2.13197 | 0.0259429 | -2.73977 | 0.0097778 |
| BCR | -1.88744 | 0.0169552 | -1.69111 | 0.0063571 | -1.92401 | 0.0224 | -2.6232 | 0.00775 |
| CTPS2 | 2.11481 | 0.0164667 | 2.02244 | 0.009410 | 1.9725 | 0.034264 | 1.40847 | 0.038 |
| KCNJ16 | 3.07469 | <0.000001 | 3.18708 | <0.000001 | 2.70281 | 0.0258667 | 3.10429 | 0.0047273 |
| PRKA | -1.14153 | 0.0410331 | -1.7829 | 0.0097886 | -1.21833 | 0.0392353 | -1.3353 | 0.0218218 |
| SPATA5 | -3.47076 | <0.000001 | -3.37156 | <0.000001 | -2.99732 | 0.0266667 | -3.82648 | 0.0074286 |
| TMEM256 | 2.29439 | 0.0270083 | 3.48716 | <0.000001 | 3.3222 | 0.014 | 2.26091 | 0.0429375 |

**Fig. S9. Quantitative proteomic analysis of renal stress-resistance mediated by tailored dietary interventions in kidneys.**

**(A)** Principal component analysis of all dietary conditions examined in a single proteomic dataset. **(B)** Overlap of differentially regulated renal proteins after dietary preconditioning with FMD, SR80/SR100 and CR as compared to non-preconditioned animals ($p<0.05$). **(C)**. $Log_2$ Fold Change and q-values for overlapping, differentially regulated proteins in response to FMD, SR80/100 and CR compared to the non-PC group, respectively. n = 4 biological replicates per dietary group. Abbreviations: BCAA: Dietary Restriction of Branched Chain Amino Acids, CR: Caloric Restriction; FMD: Fasting Mimicking Diet; KD: Ketogenic Diet;

non-PC: non-preconditioned mice; SR100: 100% Restriction of Sulfur-containing Amino Acids, SR80: 80% Reduction of Sulfur-containing Amino Acids.

| Weight loss | Compared to baseline prior renal ischemia-reperfusion injury |
|---|---|
| 0 points | no weight loss |
| 1 point | < 5% |
| 5 points | 5 - 10% |
| 10 points | 11-19% |
| 20 points | 20% |

| Appearance | |
|---|---|
| 0 points | smooth fur, clean body-openings, clear and shiny eyes |
| 1 point | shaggy fur, dull skin and/or scaly skin, hair loss without skin pathologies, small excoriations (superficial and locally circumscribed without rush or swelling) |
| 10 points | clotty fur and/or clotty body-openings, dull eyes, head tilting, body tilting, moderate excoriations/skin pathologies (with rush or swelling), impaired wound healing |
| 20 points | severe excoriations or skin pathologies (severe rush, swelling, purulent discharge, no wound-healing), abscess, tense abdomen, distended abdomen, paralysis) |

| Activity | |
|---|---|
| 0 points | normal appearance (flight reflex, nest building, fur cleaning, movement in cage) |
| 5 points | decelerated movement and/or occasional stereotypies |
| 10 points | self-isolation, occasional lethargy with half-closed eyes, severe hyperactivity, severe stereotypies (e.g. spinning-around) and/or impairment of coordination (e.g. staggering and/or reeling) |
| 20 points | self-aggression/ -mutilation, apathy (fully closed eyes) |

| Total sum | |
|---|---|
| < 10 points | continue experiment according to protocol |
| 11-19 points | close observation and consultation with experimental supervisor |
| > 20 points | immediate sacrifice of the animal |

**Table S1: Recovery score after renal ischemia-perfusion injury (IRI). A recovery score based on weight loss, appearance and activity was examined daily as a read-out for postoperative heath status after renal IRI in rodents.**

The recovery score was compiled based on the “Belastungskataloge zur Bewertung von Tierversuchen” of the „Forum Tierversuche in der Forschung“ and approved by the Landesamt für Natur, Umwelt und Verbraucherschutz Nordrhein-Westfalen (State Agency for Nature,

Environment and Consumer Protection North Rhine-Westphalia). Abbreviation: IRI: Ischemia Reperfusion Injury.

| | CR | | FMD | | SR100 | | SR80 | |
|---|---|---|---|---|---|---|---|---|
| Protein | $Log_2FC$ | q-value | $Log_2FC$ | q-value | $Log_2FC$ | q-value | $Log_2FC$ | q-value |
| CBS | -0.23261 | 0.55609 | -0.32796 | 0.38007 | -0.62064 | 0.03212 | -0.56219 | 0.13339 |
| CDO1 | 0 | 1.00000 | 3.42248 | 0 | 0 | 1.00000 | 0 | 1.00000 |
| CSAD | -0.39069 | 0.20124 | -0.41000 | 0.18281 | -0.95686 | 0.00651 | -0.00165 | 0.99789 |
| CTH | 0.20199 | 0.39012 | -0.16163 | 0.47749 | 0.02574 | 0.90141 | -0.38309 | 0.25493 |
| ETHE1 | -0.12724 | 0.65480 | 0.11173 | 0.67948 | -0.11101 | 0.78495 | 0.19846 | 0.64087 |
| GCLC | -0.07067 | 0.86667 | -0.04767 | 0.88741 | 0.41800 | 0.14199 | 0.49071 | 0.13429 |
| GCLM | -0.37074 | 0.27947 | 0.28284 | 0.35538 | -0.18548 | 0.48961 | 0.29001 | 0.48856 |
| GOT1 | 0.14320 | 0.62325 | 0.36232 | 0.13688 | 0.33562 | 0.09496 | 0.05991 | 0.83547 |
| GOT2 | 0.17816 | 0.57828 | 0.43069 | 0.12689 | 0.26818 | 0.27852 | 0.17092 | 0.61737 |
| GSS | -0.19898 | 0.52379 | 0.11060 | 0.65603 | -0.60364 | 0.03009 | -0.24403 | 0.47154 |
| MPST | 0.50365 | 0.12483 | -0.03119 | 0.92434 | 0.76458 | 0.01341 | -0.07888 | 0.87000 |
| SUOX | 0.18499 | 0.57592 | 0.25605 | 0.40033 | 0.36938 | 0.17829 | 0.32322 | 0.44870 |
| SRQDL | -0.10803 | 0.76662 | 0.05346 | 0.87694 | -0.30599 | 0.20391 | 0.01981 | 0.96201 |
| TST | 0.37429 | 0.23844 | 0.20324 | 0.40262 | 0.40088 | 0.08583 | 0.13633 | 0.63494 |

**Table S2: Changes in proteomic expression pattern of key enzymes in oxidative and hydrogen sulfide ($H_2S$)-dependent cysteine catabolism in murine kidneys after FMD, SR80/100 and CR.**

Depicted are $\log_2$ Fold Change and q-values for key enzymes of oxidative and $H_2S$-dependent cysteine catabolism as established by proteomic analysis of kidneys of dietary preconditioned compared to non-preconditioned animals. Abbreviations: CR: Caloric Restriction; FMD: Fasting Mimicking Diet; SR100: 100% Restriction of Sulfur-containing Amino Acids; SR80: 80% Reduction of Sulfur-containing Amino Acids.

| Variables<br>Included cases | Control<br>10 | Low-SAA<br>10 | p-value |
|---|---|---|---|
| Age (years), median (IQR) | 58 (56-62) | 58 (54-62) | 0.796 |
| Male, n (%) | 5 (50.0) | 5 (50.0) | 1.000 |

**Table S3: Baseline demographics of participants in the pilot-cohort adhering to control or low-SAA diet**.

Age is depicted as median and interquartile range (IQR). P-values were calculated by Mann-Whitney-U-Test in numerical and by $\chi^2$-test in categorical variables, respectively.

| Item number | FMD Day 1 S5159-E736 | FMD Day 2-3 S5159-E738 |
|---|---|---|
| **Proximate Content** | | |
| Broth mix (g) | 5.95 | 5.31 |
| EVOO (g) | 12.75 | 0.0 |
| Essential fatty acid (g) | 0.21 | 0.0 |
| Vegetable mix (g) | 14.88 | 0.0 |
| Glycerol (g) | 0.0 | 0.75 |
| Hydrogel | 56.7 | 94.96 |
| Fiber (g) | 5.0 | 0.0 |
| Mineral (AIN-93G-MX), Teklad, TD 94046, ENVIGO | 3.5 | 0.0 |
| Vitamin (AIN-93-VX), Teklad, TD 94047, ENVIGO | 1.0 | 0.0 |
| **Metabolizable energy [MJ/kg][1]** | 7.67 | 1.48 |
| kcal Protein (%) | 5.0 | 0.5 |
| kcal Fat (%) | 28.0 | 0.5 |
| kcal Carbohydrates (%) | 64.0 | 99.0 |

**Table S4: Proximate content and metabolizable energy of the day 1 and day 2-3 components of the FMD.**

FMD consist of two different dietary components designated as day 1 and day 2-3 diet, respectively. After the end of FMD, standard chow (Table S4) was fed ad libitum for 4 days. FMD was produced by ssniff Spezialdiäten GmbH, Soest, Germany as previously described (*13*). Specified item numbers are depicted. [1] Metabolizable energy was calculated according to the Atwater formula corresponding to 3.275 kcal/kg. Abbreviations: EVOO: Extra Virgin Olive Oil; FMD: Fasting Mimicking Diet; kcal: kilocalories.

| Item number | KD<br>S51519-E730 | BCAA<br>S5159-E724 | SR100<br>S5159-E720 | SR80<br>S5159-E722 | Standard chow<br>V1554-330 |
|---|---|---|---|---|---|
| **Proximate content** | | | | | |
| Crude protein (%) | 15.9 | 19.5 | 11.9 | 11.9 | 19.3 |
| Crude fat (%) | 63.1 | 8.1 | 4.5 | 4.5 | 3.3 |
| Crude fiber (%) | 0.0 | 3.0 | 4.9 | 4.9 | 4.4 |
| Crude ash (%) | 6.0 | 3.9 | 3.2 | 3.2 | 6.0 |
| Starch (%) | 0.3 | 15.8 | 51.2 | 51.2 | 36.0 |
| Sugar (%) | 0.0 | 29.6 | 19.9 | 19.9 | 5.0 |
| Dextrin (%) | 0.0 | 16.2 | 0.0 | 0.0 | 0.0 |
| Lysine (%) | 1.33 | 1.59 | 1.09 | 1.09 | 1.04 |
| Methionine (%) | 0.76 | 0.66 | 0.0 | 0.09 | 0.37 |
| Cysteine (%) | 0.07 | 0.71 | 0.0 | 0.07 | 0.37 |
| Valine (%) | 1.11 | 0.27 | 0.79 | 0.79 | 1.04 |
| Isoleucine (%) | 0.90 | 0.25 | 0.79 | 0.79 | 0.84 |
| Leucine (%) | 1.58 | 0.82 | 1.07 | 1.07 | 1.64 |
| Glutamic acid (%) | 3.59 | 2.87 | 3.50 | 3.32 | 3.68 |
| Calcium (%) | 0.89 | 0.63 | 0.52 | 0.52 | 1.0 |
| Phosphorus (%) | 0.52 | 0.48 | 0.34 | 0.34 | 0.7 |
| Sodium (%) | 0.45 | 0.11 | 0.15 | 0.15 | 0.23 |
| Magnesium (%) | 0.25 | 0.05 | 0.07 | 0.07 | 0.22 |
| Potassium (%) | 1.63 | 0.37 | 0.48 | 0.48 | 0.73 |
| **Metabolizable energy [MJ/kg][1]** | 26.6 | 17.2 | 16.0 | 16.0 | 16.4 |
| kcal Protein (%) | 10.0 | 19.0 | 12.0 | 12.0 | 23.0 |
| kcal Fat (%) | 89.0 | 18.0 | 11.0 | 11.0 | 10.0 |
| kcal Carbohydrates (%) | 0.6 | 63.0 | 77.0 | 77.0 | 67.0 |

**Table S5: Proximate content and metabolizable energy of KD, BCAA, SR80/100 and the standard chow used for dietary preconditioning.** All diets were obtained from ssniff Spezialdiäten GmbH, Soest, Germany and specified item numbers are depicted. [1] Metabolizable energy was calculated according to the Atwater formula corresponding to 3.275 kcal/kg. Abbreviations: BCAA: Dietary Restriction of Branched Chain Amino Acids; kcal: kilocalories; KD: Ketogenic Diet; SR100: 100% Reduction of Sulfur-containing Amino Acids; SR80: 80% Reduction of Sulfur-containing Amino Acid
